# Role of Divalent Ions in Membrane Models of Polymyxin-Sensitive and Resistant Gram-Negative Bacteria

**DOI:** 10.1101/2024.08.29.610330

**Authors:** Mariia Savenko, Robert Vácha, Christophe Ramseyer, Timothée Rivel

## Abstract

Polymyxins, critical last-resort antibiotics, impact the distribution of membrane-bound divalent cations in the outer membrane of Gram-negative bacteria. We employed atomistic molecular dynamics simulations to model the effect of displacing these ions. Two polymyxin-sensitive and two polymyxin-resistant models of the outer membrane of Salmonella enterica were investigated. First, we found that the removal of all calcium ions induces a global stress on the model membranes, leading to substantial membrane restructuring. Next, we used enhanced sampling methods to explore the effects of localized stress by displacing membrane-bound ions. Our findings indicate that creating defects in the membrane-bound ion network facilitates polymyxin permeation. Additionally, our study of polymyxin-resistant mutations revealed that divalent ions in resistant model membranes are less likely to be displaced, potentially contributing to the increased resistance associated with these mutations. Lastly, we compared results from all-atom molecular dynamics simulations with coarse-grained simulations, demonstrating that the choice of force field significantly influences the behavior of membrane-bound ions under stress.

## INTRODUCTION

The growing occurrence of multidrug-resistant (MDR) Gram-negative bacteria poses a significant threat to public health^1–3^. Of particular concern is the resistance to last-resort antibiotics, such as those in the polymyxin family, which are crucial for treating infections caused by these resilient pathogens. Despite extensive efforts from the scientific and political institutions to develop novel antibiotic therapies, the approval rate for new drugs has been alarmingly low in recent years^4–6^. This trend is exacerbated by a limited understanding of the dynamic changes within the cell envelope of Gram-negative bacteria under environmental stresses.

The Gram-negative cell envelope, comprising an inner membrane (IM) and an outer membrane (OM) separated by a peptidoglycan layer, plays a fundamental role in protecting bacteria from external threats, including antibiotics. The OM, with its unique asymmetric structure featuring phospholipids on the inner leaflet and lipopolysaccharides (LPS) on the outer leaflet, significantly contributes to the low permeability of Gram-negative bacteria to drugs^7–9^. The presence of integral proteins enriches the functions of this membrane^10,11^ and plays an important role in the modes of action of some antibiotics.

In this work, we focus on the interactions of antibiotics with the lipid matrix of the OM, particularly with lipopolysaccharides. LPS are amphiphilic molecules that expose a substantial hydrophilic layer, including O-antigen and parts of core oligosaccharides^12–14^. This hydrophilic layer interacts with the external environment, and which is crucial for bacterial survival under stress conditions such as antibiotic exposure^15^. Modifications in LPS structure enable bacterial resilience to antibiotics, contributing to the diverse chemotypes and adaptive resistance mechanisms observed in Gram-negative bacteria^16,17^.

Among the limited arsenal of effective antibiotics against MDR Gram-negative bacteria, polymyxins B (PMB) and E (PME, colistin) stand as last-resort treatments^18–21^. Initially relegated due to their nephro- and neurotoxic effects^22,23^, polymyxins have regained prominence in combating antibiotic-resistant infections^15,24^. Their unique structure, consisting of a cyclic heptapeptide linked to a fatty acid tail by a central tripeptide, facilitates specific interactions with the OM, particularly by means of their positively charged L-diaminobutyric acid (DAB) residue. The cyclic peptide structure is also critical for their antibacterial activity, distinguishing them from less effective linear analogs^21,25,26^.

The mechanism by which polymyxins exert their bactericidal effect involves multiple stages of interaction with the OM^13,15,27–31^, some of which remain under debate. Upon contact, polymyxins bind to the OM surface, facilitated by their cationic nature. Polymyxins accumulate in regions of membrane defects^27,32^, *i.e*. regions involving local modifications in membrane structure, although the exact nature of these defects and their impact on polymyxin action when adsorbed to the membrane are still unclear. Previous works have indicated that polymyxin B may aggregate in solution^33^ and on the surface of OM models, forming micelle-like structures^27,28,32^. However, it is unclear whether polymyxin aggregation is necessary for effective bactericidal effect.

Following adsorption to the OM, polymyxins are believed to superficially insert their fatty acid chain into the membrane^21^. This partial insertion is presumably made possible by the combined presence of the fatty acid chain^25^, the cyclic peptide structure, and the presence of the cationic DAB residues. It is believed that these residues displace membrane-bound divalent cations^15,18,20,21,23,34^, *i.e*. Ca^2+^ or Mg^2+^. This displacement weakens the OM structure, potentially allowing polymyxins to penetrate deeper into the lipid bilayer and exert their antibacterial effects.

However, bacterial resistance to polymyxins has emerged as a growing concern, necessitating a deeper understanding of resistance mechanisms employed by Gram-negative bacteria. Resistance often involves alterations to LPS, such as the addition of chemical groups like phosphoethanolamine (PEtN)^31,35–37^, 4-aminoarabinose (Ara-4N)^31,35,36^, or galactosamine (GalN)^38^, which tends to neutralize the overall negative charge of LPS^39^ and diminish repulsive interactions with polymyxins, which subsequently rely less on the presence of divalent ions. Understanding the implications of these resistance mechanisms on the interactions between polymyxins and the OM is critical for developing strategies to combat polymyxin resistance.

To elucidate these intricate processes, molecular dynamics (MD) simulations provide a powerful tool for studying atomic-scale interactions and dynamics over sub-millisecond timescales. Recent studies have highlighted conflicting findings regarding the influence of polymyxins on membrane-bound divalent cations^27,40–43^, emphasizing the need for further investigation into their precise mode of action and resistance mechanisms.

In this work, we aim to address these gaps by employing a computational approach to investigate the consequences of divalent ion displacement on OM properties and its implications for polymyxin efficacy. Through all-atom simulations, we model a global stress involving the removal of all divalent ions in both colistin-susceptible and colistin-resistant model membranes of *Salmonella enterica*. Additionally, we introduce a novel collective variable to model a local stress on the divalent ions that could be induced by polymyxins. Finally, we applied this collective variable to coarse-grained simulations which can reach longer timescales.

## RESULTS

### Reaction of outer membrane to removal of divalent ions

Divalent ions, particularly Ca^2+^ and Mg^2+^, play a bridging role between lipopolysaccharides of the outer membrane of Gram-negative bacteria. To better understand the impact of their displacement on membrane properties, we first modeled the global removal of these ions for two polymyxin-sensitive and two polymyxin-resistant models. Figure 1 shows that global stress creates major membrane remodeling, including LPS flipping as well as cracks at the surface of LPS layer, which are openings in the membrane that go through a whole monolayer, in the polymyxin-sensitive model membrane referred to as P2.

**Figure 1.**
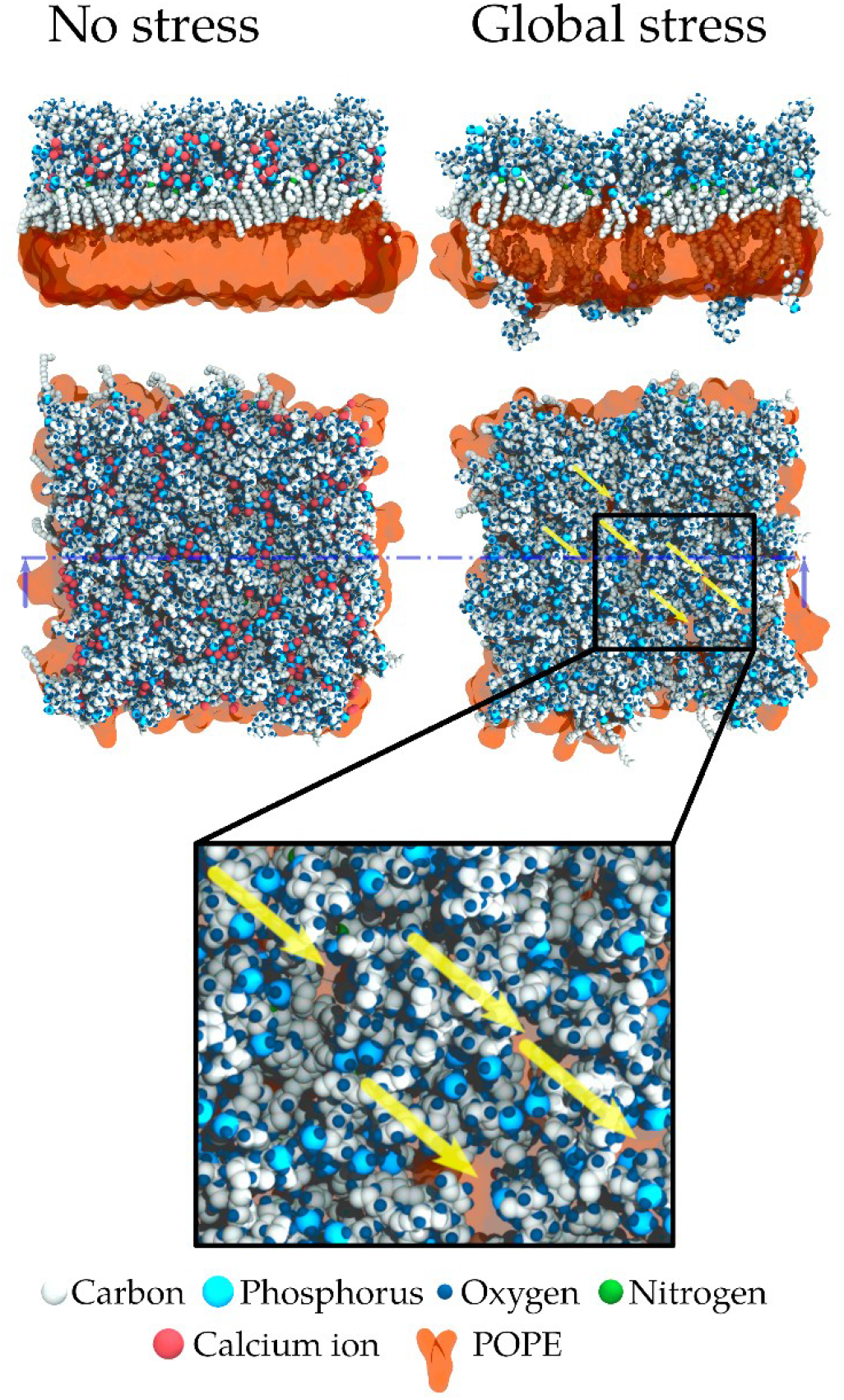
Representative snapshots from our simulations of one model of outer membrane of Gram-negative bacteria (system P2), in the absence of stress and under a global stress originating from the removal of all calcium ions in the box (and their replacement by sodium in the solvent). Global stress induces the formation of cracks in each leaflet. Yellow arrows show some of these cracks that reside on the LPS leaflet.

The signature of calcium ions removal is clearly recognizable in **Figure 1**, in our simulations relying on CHARMM36 force field^44^. Indeed, on the side view of the system with no stress applied, a first layer of phosphorus atoms and calcium ions, associated to the lipid A phosphate groups, and a second larger layer, associated to LPS inner core phosphate groups are recognizable (see the density profile in Figure S 1 as well), while this structure is clearly hampered in the system where all divalent ions were removed, and where the membrane profile looks much rougher (see also Figure S 2 and S 5).

We compared the effect of the global stress in four different membrane models (see **Figure 2**). The first model, P2, is the canonical model of a sensible strain of *Salmonella enterica*. A variant of that model, P1, assigns a net charge of -1 to the lipid A phosphate groups, as recommended by Rice et al^45^ Next two models are associated to mutations in the lipopolysaccharides held responsible to polymyxin resistance. The first of them, PEtN, binds phosphatidylethanolamine group to the lipid A phosphate groups, while the second, Ara-4N, decorates them with 4-amino-4-deoxy-l-arabinose. This set of model membranes allows us to compare their susceptibility to sustain stress and its impact on the content and distribution of divalent ions.

**Figure 2.**
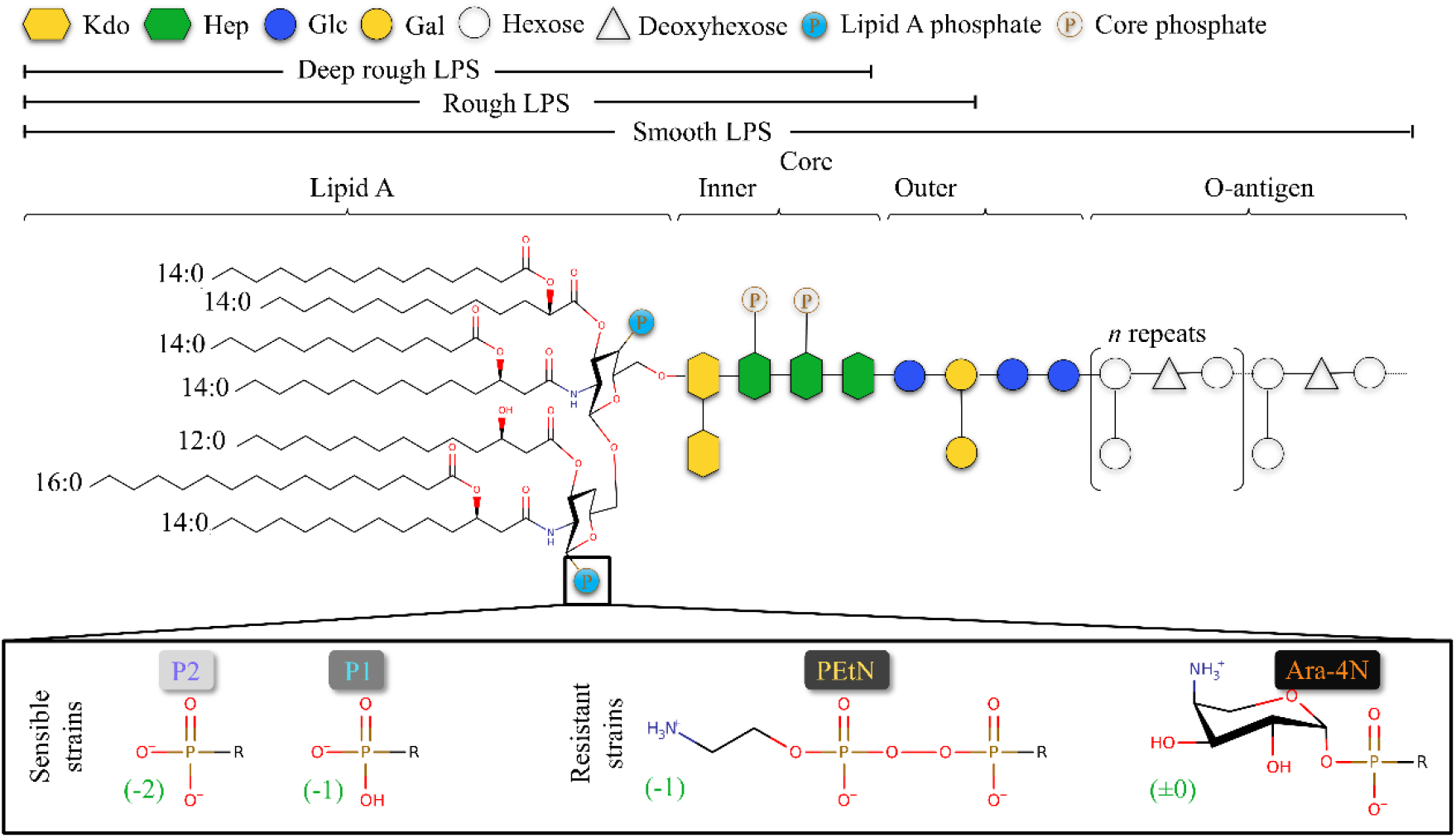
Chemical structure of the four models of LPS employed. The variation resides in the phosphate groups attached to the lipid A (box at the bottom) with two models featuring phosphate groups that have different net charges (−2 or -1, depending on the protonation state). The other two models are decorated with PEtN or Ara-4N groups, which are associated with polymyxin resistance in bacterial strains.

In this regard, we compared the packing defect (**Figure 3**) induced by global stress on the four model membranes. We generated four replicas for each system all of which were simulated until the variations in the density profiles of different relevant groups do not change with time after 300 ns (see Figure S 2 – S 5) and we calculated the defect size. The removal of all calcium ions in the system drastically affects all model membranes that we studied. We observe a multifold increase of the deep defect size for all model membranes. Even in the case of PEtN, where the characteristic size of deep defects increases the least, it still grows from 7.0 ± 0.1 Å to 17 ± 4 Å. On the other hand, we see that P2 is the system that is most affected by such stress for both types of packing defects.

**Figure 3.**
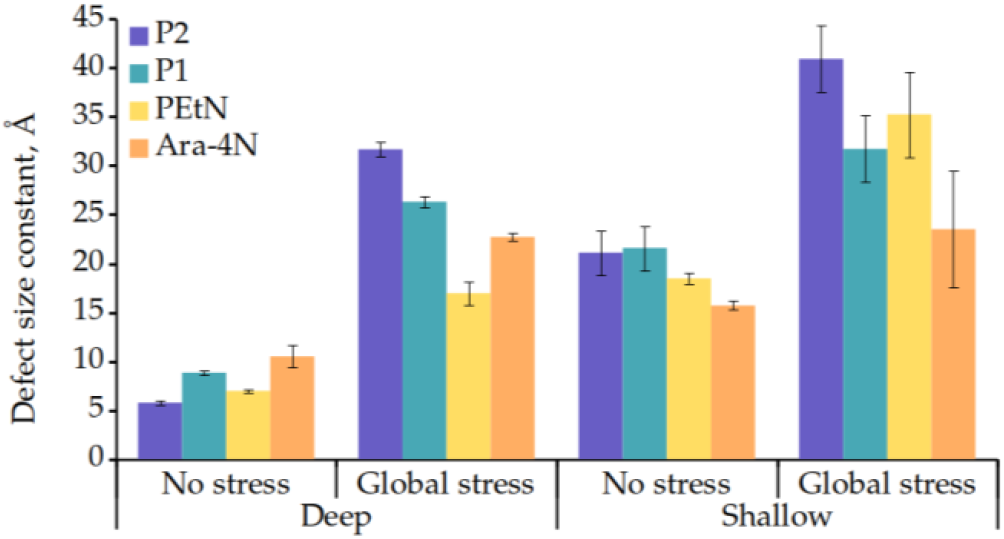
Defect size constant for equilibrated membranes and membranes under global stress

In the presence of calcium ions, the density of water molecules decreases more rapidly along the normal to the membrane, indicating that water penetrates the membrane less deeply. Susceptible strains (P2 and P1 models) display an increase in water in the surrounding of both layers of phosphorus / calcium ions which fades when the ions are removed. Removal of divalent ions drives water much deeper, presumably through the cracks that are observed in the positions of LPS flipping (see Figure S 6).

We calculated the distribution of the area per lipid. In the case of the system without stress (Figure S 7 – S 8) LPS area per lipid for both P2 and P1, share the same value (1.93 ± 0.01). We also notice that the largest area per lipid is for PEtN with a value of 2.11 ± 0.02, while Ara-4N displays a slightly lower area per lipid (2.07 ± 0.03). The removal of calcium ions dramatically widens the area per lipid distribution (Figure S 7). Interestingly, the widening is most significant for P2, followed by P1, PEtN, and Ara-4N. However, it is not straightforward to estimate accurately the average area per lipid in systems that suffered such major stress, because of the drastic widening of the distribution. Indeed, with the presence of cracks on the LPS monolayer, it is questionable whether the calculation of the area per lipid should exclude them, rendering the calculation substantially a harder task.

To complement the insight obtained with the area per lipid distribution, we also investigated the proportion of LPS that flipped through the simulation, as this observable shall be directly related to the change in LPS leaflet area. Table 1 shows the proportion of LPS that flipped from upper to lower leaflet.

**Table 1.** Percentage of LPS that flipped towards the lower leaflet (POPE), and the standard error associated. The proportion is computed from the density profile of the head group as reported in the supplementary information (see Figure S 9 – S 12).

| System | Proportion of flipped LPS | Error estimate |
| --- | --- | --- |
| P2 | 11 % | 2 % |
| P1 | 9 % | 2 % |
| PEtN | 5 % | 1 % |
| Ara-4N | 1.3 % | 0.6 % |

### Modeling local ion displacement

In the previous section, we addressed an extreme case in which the global stress associated with complete ions removal could be referred to as a detergent effect. On the other hand, that did not cover the local mechanism of ion displacement and its effects on membrane properties. In this section we introduce a new collective variable that aims at describing the local polymyxin-induced ion displacement. Polymyxins, being polycationic lipopeptides, are believed to displace calcium or magnesium ions by means of repulsive electrostatic interactions. A straightforward way to model ion displacement would be to move those ions radially from the position of a defect that is considered bound to one or several polymyxins. It should be noted that such a model does not require the presence of any polymyxins and focuses only on the influence of ion displacement and thus, does not involve direct interactions between polymyxins and membrane. We propose a new collective variable ξ_dis_, that controls the number of calcium ions in a cylinder spanning transversally to the membrane.

Figure 4. shows density maps and representative snapshots from MD simulations of the calcium ions in system P2, for different values of the collective variable, where ξ_dis_ is expressed in number of ions per area unit, in the cylinder. This variable allows us to control the number of calcium ions in a local portion of the membrane (see also Movie S1 for the action of the collective variable on the calcium ions during a steered MD simulation). Interestingly, we do not see major membrane remodeling upon such local stress, as it is depicted both in and **Figure 4** and **Figure 5**. Our results show a significant increase in the deep packing defect constant only in the case of P2 (see Figure S 13).

**Figure 4.**
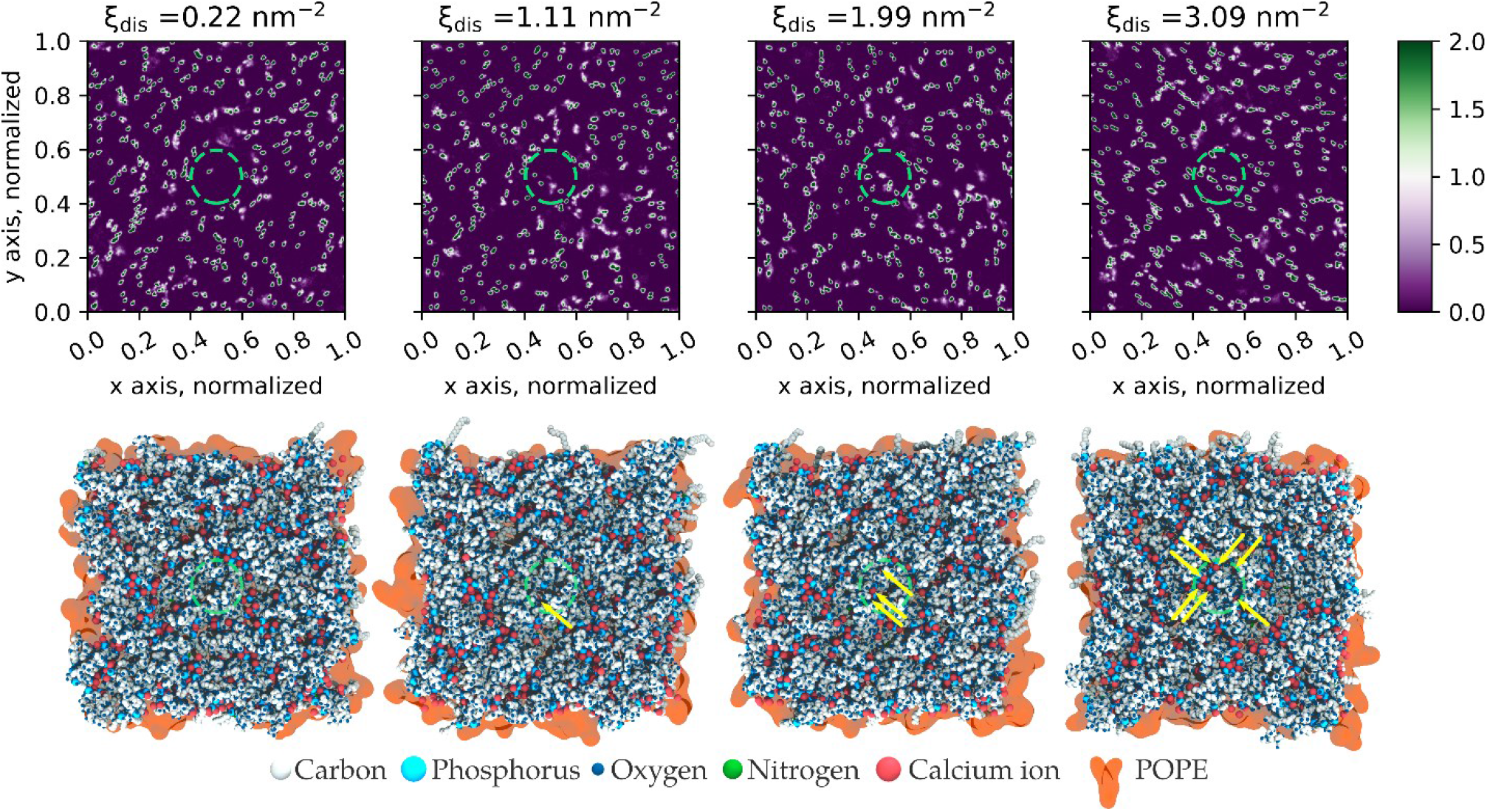
The upper panels show the density maps of calcium ions for a given value of the collective variable. Each map is associated with a representative snapshot of the system. Arrows point at calcium ions that are visible directly from the top view. The circles (in green) represent the cylinder where the biasing potential is applied. Number density values are in nm^-3^.

**Figure 5.**
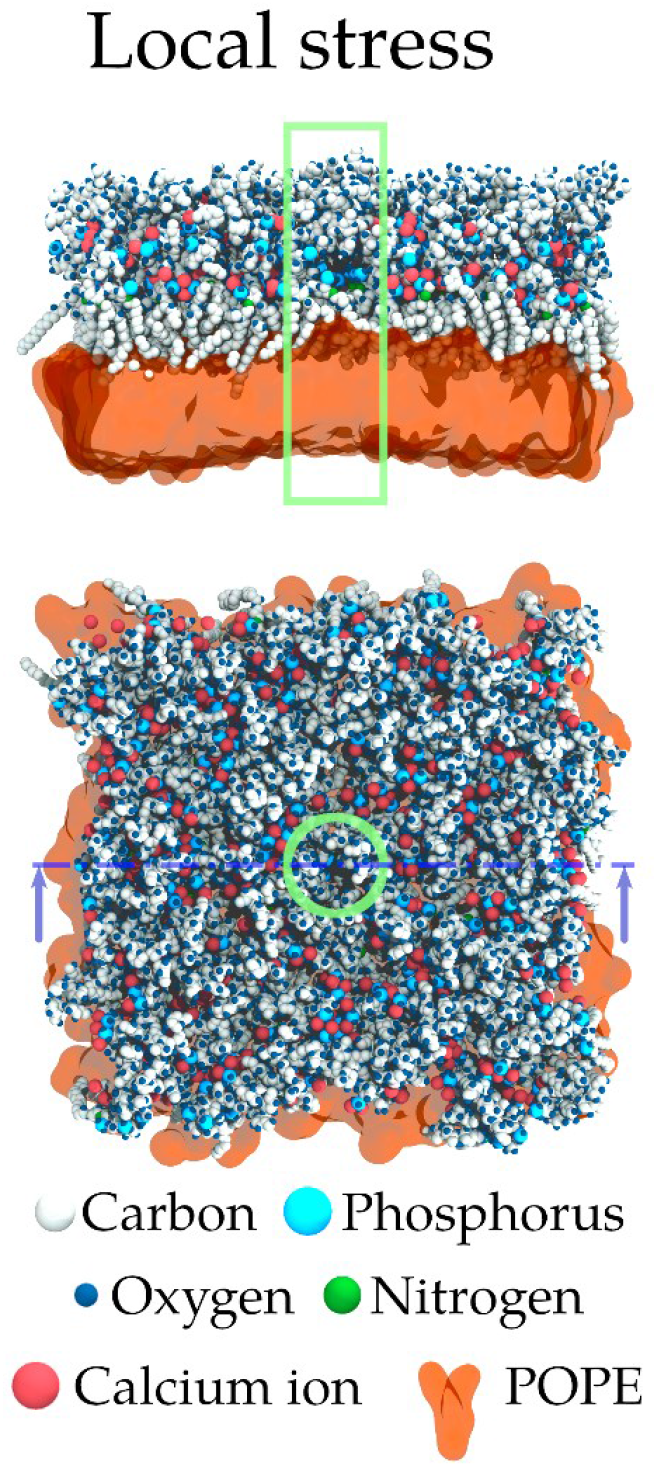
Representative snapshot from our simulations of one model of outer membrane of Gram-negative bacteria (system P2) under a local stress consisting in the local displacement of calcium ions away from a cylindrical volume centered in the simulation box (in green).

Figure 6. shows the variation in the number of phosphorus atoms in the cylinder with the values of the collective variable. P2, P1, and Ara-4N exhibit remarkably similar trends in the variation of the number of phosphates, which seems linearly correlated to the collective variable. The discrepancy observed for PEtN can be explained by the presence of one extra phosphorus atom in each of the lipid A phosphate group. This effect of co-motion of the phosphorus atoms does not mean that the whole LPS molecules are moving away from the cylinder. A small portion of the LPS, close to the phosphate groups, seems to leave the cylinder, while some LPS molecules rotate to ensure that the phosphorus atoms interacting with the displaced calcium ion(s) preserve this interaction (see also Movie S2 that shows the co-motion of phosphorus atoms along with biased calcium ions). This collective motion P-Ca^2+^ is a marked difference between local and global stress, which could play a role in the global conservation of membrane properties in case of such local stress, by avoiding direct repulsion between unbridged phosphate groups.

**Figure 6.**
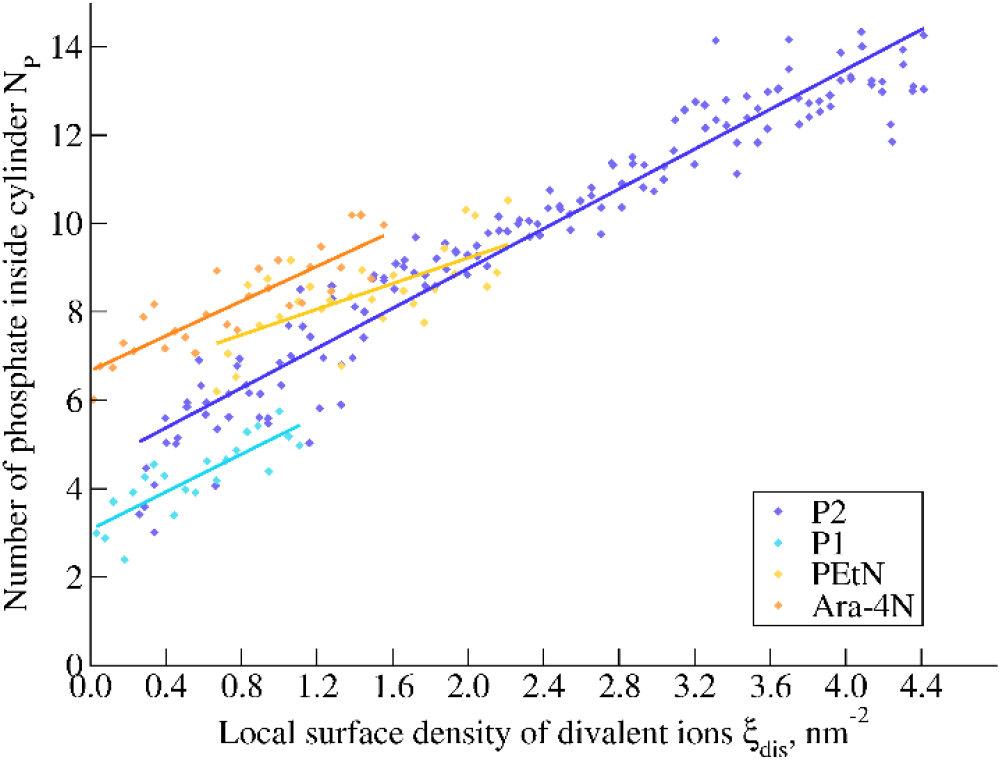
Variation of the number of phosphorus atoms in the cylinder as a function of the collective variable. Linear regressions are also plotted.

Note that the local stress has small effects on global membrane properties. For instance, there is no representative impact on the distance between phosphorus atoms of different LPS molecules (see Figure S 14), nor any trends in the orientation of lipid A head groups (see Figure S 15). This stress has also little effect on membrane dynamic properties (see Figure S 16). However, the local stress widened the area per lipid distribution, especially for P2, PEtN, and Ara-4N (see Figure S 8), while changes are not significant in the case of P1.

We computed the free energy profile of calcium displacement, along our collective variable (see Methods for further details), see Figure 7. To better estimate the surface density of calcium ions in each membrane in the absence of stress, we fitted a quadratic function that reads 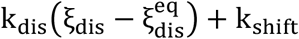 to the free energy profiles, in the close vicinity to the equilibrium value. P2 has the highest minimum at 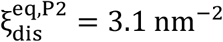, while both P1 and Ara-4N share equilibrium value, at 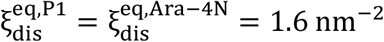. The equilibrium value for PEtN stands in between, at 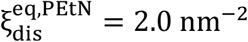. This points out that the net charge of LPS phosphate groups is not the only contribution to the divalent ions content in the membrane. We also estimated the average value 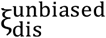 for unbiased equilibrated membrane models, see **Table 2**. The differences observed in **Table 2**, between 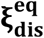 and 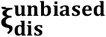 suggest that the calcium density is not in equilibrium and is rather dictated by the membrane generator implemented in CHARMM-GUI where the number of calcium ions inserted is based on the charge of LPS phosphate groups.

**Table 2.** Comparison of the equilibrium surface density of calcium ions from fitted free energy minima and unbiased membrane simulations.

| System | $\xi_{\text{dis}}^{\text{eq}}, \text{nm}^{-2}$ | $\xi_{\text{dis}}^{\text{unbiased}}, \text{nm}^{-2}$ |
| --- | --- | --- |
| P2 | 3.1 | 2.6 |
| P1 | 1.6 | 1.4 |
| PEtN | 2.0 | 1.9 |
| Ara-4N | 1.6 | 1.8 |

**Figure 7.**
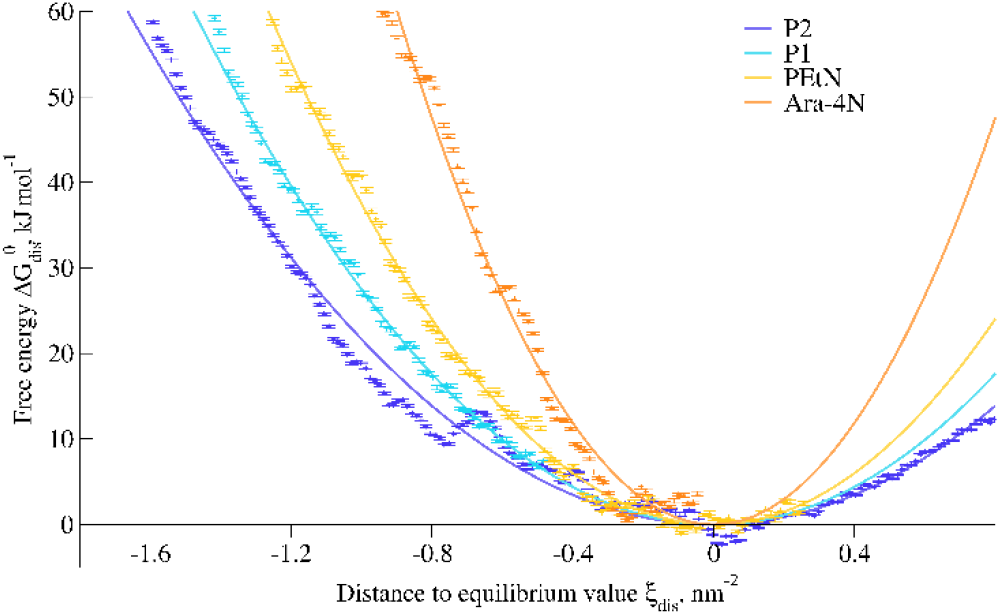
Free energy profiles of calcium displacement for various model membranes as a function of our collective variable. The profiles are shifted so that the minimum value of their quadratic fit is set to zero.

The free energy profiles allow us to better understand the contributions of the mutations affecting lipid A phosphate groups. The steeper the free energy profile is, the harder it is to create a defect in the membrane-bound ion network. Therefore, the quadratic fits of these free local profiles allow us to quantify the response of the system to a local stress. In this case, we can sort the different system based on the coefficient k_dis_, and see that 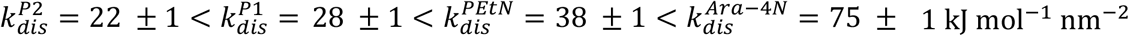. Note that lower values of k_dis_ correlate with higher propensity of flipping in the case of global stress.

### Local stress and the presence of polymyxin

We simulated P2-PME system, that contains one polymyxin E or colistin, restrained to the region of the cylinder. Since we are interested in the early stages of polymyxin modes of action, we decided to place the polymyxin molecule close to the top of LPS layer. We want to address first, whether polymyxin assist the formation of the defect by making divalent ion displacement less energetically costly, and secondly whether the presence of such a defect assist polymyxin permeation through the outer membrane.

Figure 8 shows the free energy profile of calcium displacement for both P2 and P2-PME. There is no significant variation of the equilibrium value of the local surface density of divalent ions, as 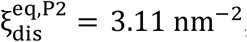, while 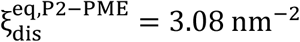, which can be explained by the presence of only a single polymyxin. However, polymyxin seems to decrease the stiffness of that free energy curve, reducing the cost to modify the local ion density with 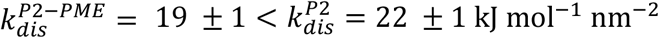.

**Figure 8.**
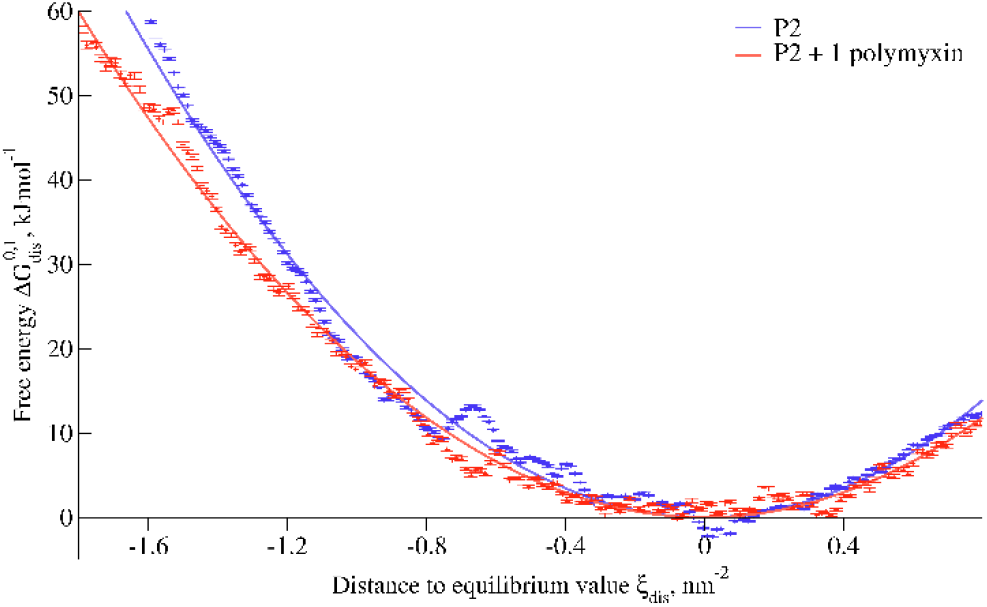
Free energy profile along our collective variable for the system P2 and P2-PME which is the model embedding one polymyxin. The profiles are shifted so that the minimum value of their quadratic fit is zero.

In case of larger stress, *i.e*. lower density of divalent ions in the cylinder, polymyxin inserts deeper in the membrane (see Figure 9). We observe a threshold around ξ_*dis*_∼1.6 nm^™2^, below which polymyxin equilibrates ∼1.0-1.5 nm deeper in the LPS leaflet.

**Figure 9.**
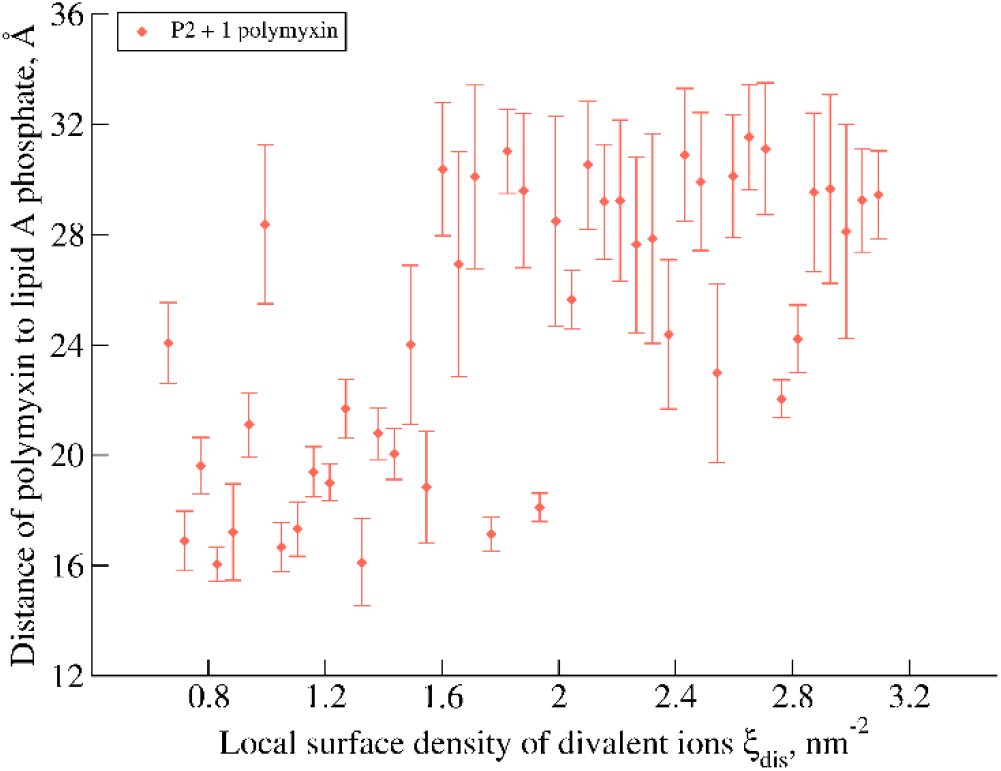
Evolution the distance of the center of geometry of polymyxin to that of lipid A phosphate groups as a function of the local surface density of divalent ions in the virtual cylinder, depicting that local stress affects the depth of insertion of polymyxin. Errors estimate refers to the standard deviation over the last 100 ns of the simulation, and the depth is the z-component of the distance of polymyxin center of geometry to lipid A phosphorus atoms.

### Local stress effects in coarse-grained simulations

Given that Martini 2 force field^44^ has been extensively used for studying interactions between polymyxins and outer membrane models^17,27,46,47^, we adapted our collective variable to this force field. At first, we computed (see Figure 10) the free energy profile with zero, one, or five polymyxin molecules (P2-CG, P2-CG-PME, and P2-CG-5PME). One can see that there is, once again, no significant variation of the equilibrium value of the local surface density of divalent ions, with 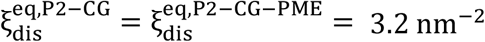. However, the addition of five polymyxin molecules drives the equilibrium value to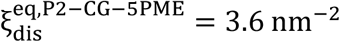, which is a surprising result as one could expect the opposite trend, *i.e*. less ions in the presence of polymyxin. It is also surprising to see that, contrary to all-atom simulations, polymyxins seem to increase the energetic cost for calcium displacement, with 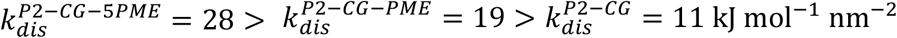.

**Figure 10.**
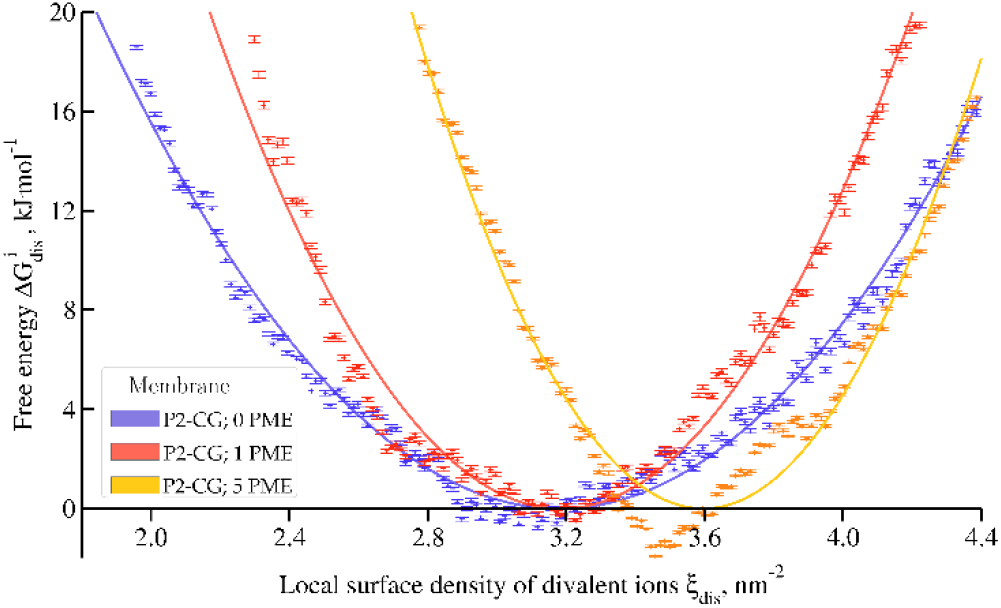
Free energy profile along our collective variable for the systems P2-CG, P2-CG-PME, and P2-CG-5PME, using coarse-grained simulations, and embedding respectively 0, 1, and 5 polymyxin molecules.

To explain the role of polymyxins in the displacement of divalent ions, we investigated the density maps of different groups in the system, namely water, LPS core, and calcium ions (see **Figure 11**). One can see that calcium beads, LPS core beads, and water present in the surrounding of lipid A head group and LPS core are *freezing* while we decrease the density in calcium in the cylinder. Freezing occurred in all coarse-grained-simulations, with and without the polymyxin molecule. It is more pronounced for lower values of collective variable, *i.e*. for lower ions densities in the cylinder.

**Figure 11.**
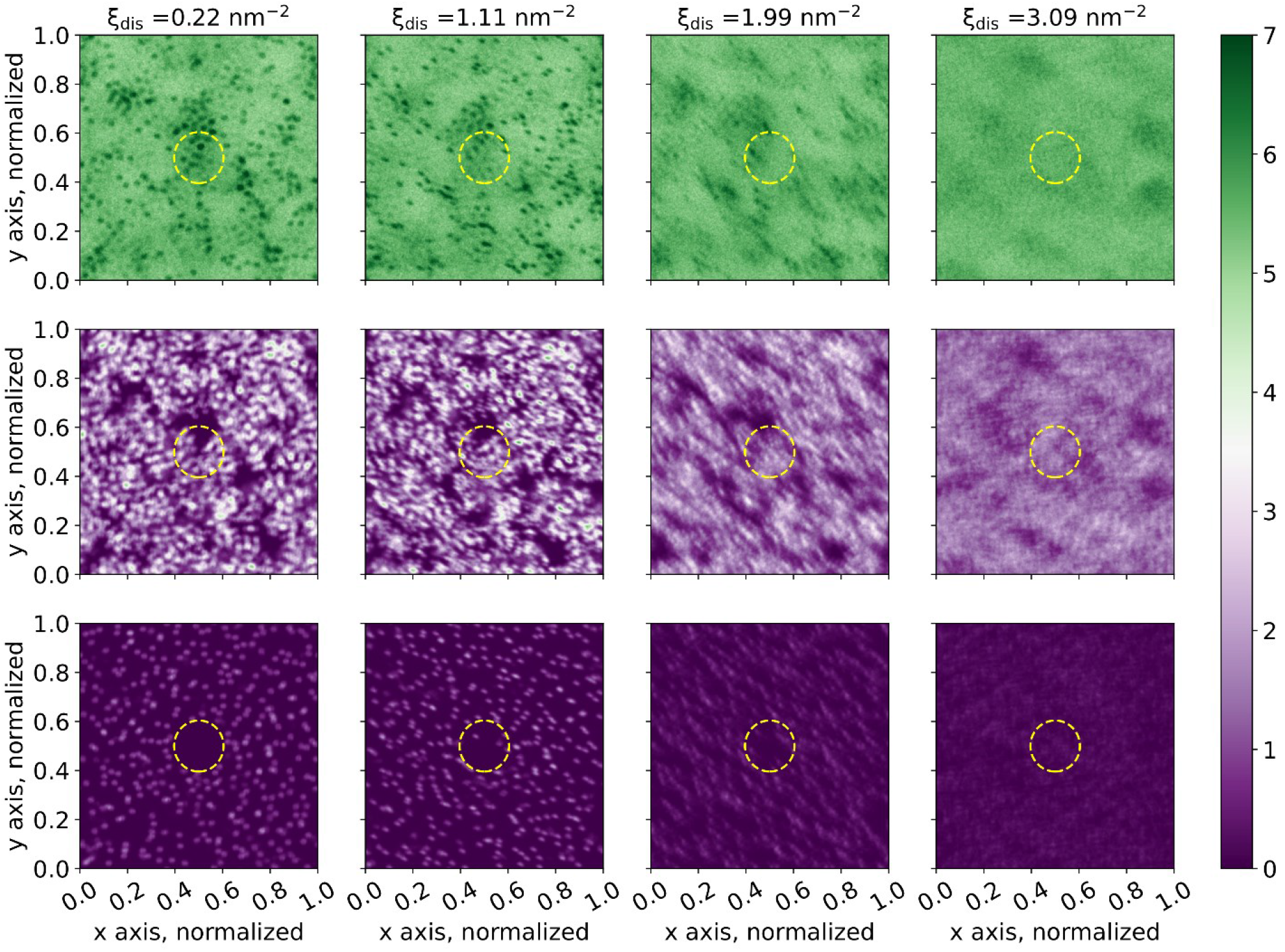
Density maps of different groups in the case of P2-CG, for four values of the collective variable. The first row shows water molecules, the second, LPS core, and the last row, shows calcium ions. Number density values are in nm^-3^. The yellow circles indicate the position of the virtual cylinder in which the biasing potential is applied.

For better understanding, we plotted the time-averaged root-mean-squared fluctuations (RMSF) of the phosphate beads that are highly affected by the freezing, as a function of the collective variable, and for different number of polymyxin molecules in the simulation box (see **Figure 12**). Such evaluation of the RMSF was already made to assess the relative flexibility of a given region of LPS^7^. We see that the more polymyxin molecules are present in the cylinder, the more phosphate beads exhibit low RMSF for higher values of the CV. We also noticed that the RMSF values drastically drop for smaller values of the collective variable, *i.e*. for low surface densities in divalent ions in the defect site.

**Figure 12.**
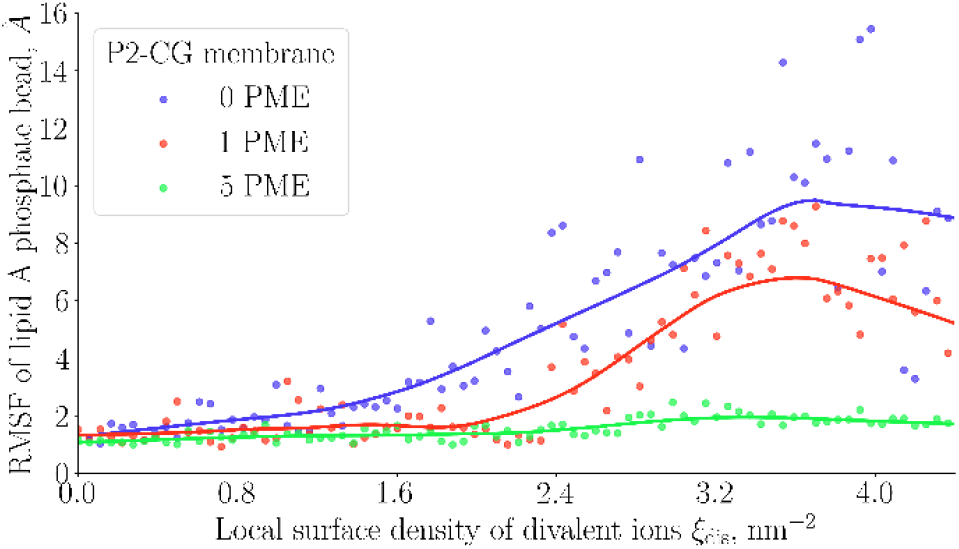
Variation of the RMSF of lipid A phosphate beads as a function of the collective variable. Plain lines are data smoothing based on local regression smoothing (LOESS).

## DISCUSSION

### The complete removal of divalent ions dramatically affects structural properties of the outer membrane

The complete loss of the calcium ions present in our model membranes induces repulsive interactions between phosphate groups of lipid A and LPS core resulting in a major and rapid increase in the upper leaflet area. The lower leaflet cannot adapt to such global stress, leading to the appearance of transient lipid islands in the lower leaflet, which does not sustain a continuous monolayer. The remodeling of the lower leaflet further drives some LPS to flip towards POPE layer, which is visible in **Figure 1**. Even though the membrane remains a bilayer in our simulations, this important remodeling could drastically affect the permeability of that membrane.

This global membrane remodeling is in agreement with works showing the chelation of membrane-bound calcium ions by ethylenediaminetetraacetic acid (EDTA)^48–51^. Cliffton and coworkers also observed membrane LPS flipping into the inner leaflet in their simulations in line with their experiments in which they wash-out calcium ions with EDTA^52^. However, it is important to consider that while comparing experimental studies using EDTA with MD simulations, where divalent ions are simply removed, can provide valuable insights, the simulations did not include the binding of EDTA into the bilayer, which may affect the results^53^. In this regard, it is interesting to note that Fu and coworkers^54^ reported that higher concentrations in polymyxin may assist lipid scrambling.

In the simulations depicted in **Figure 1**, the removal of the divalent ions did not cause a major buckling of the membrane, nor did we see an increase in the separation between leaflets as it was reported before^27^. Similarly, we do not observe loss of the characteristic lamellar structure towards an amorphous one. However, it has already been discussed that these changes in membrane structure are largely force field dependent^45,46,55^. It was reported that CHARMM36 and Martini force fields do not show such drastic changes upon divalent ions removal from the membrane, while this was observed with Gromos^55^ force field. Interestingly, GLYCAM force field was also shown to not produce stable bilayers when all cationic salts are monovalent potassium ions^45^ which may come from issues with the representation of intermolecular interactions of saccharides that tend to be overestimated^56^.

It has been commonly reported that the displacement or the (partial) removal of calcium ions in the outer membranes is a key mechanism in the action of polymyxins^28,57,58^. However, Manioglu and coworkers reported that polymyxins, upon binding to outer membrane, form crystalline structures that contribute to their antibacterial activity^59^. Furthermore, the formation of these structures is dependent on the presence of divalent ions, indicating a more complex mechanism than previously thought. While it is established that polymyxins’ interactions with outer membrane divalent ions are crucial for permeabilizing the outer membrane, it remains an open question whether polymyxins directly induce membrane rupture or if the displacement of divalent ions primarily facilitates polymyxin internalization in the bacterium. The complexity of this picture has increased as recent studies have begun to address the roles of outer membrane proteins in interactions with polymyxins^60^.

Our results, *i.e*. lipid packing defect, widening of the area per lipid distribution, and the proportion of LPS flipping, support that P2 is more sensible to divalent ions removal than polymyxin-resistant models. P1 model, in which the net charge of lipid A phosphate groups is set to -1, which represents a fully protonated state of lipid A phosphate groups^45^, seems to be an intermediate between P2 and polymyxin-resistant strains. The change in the net charge of lipid A phosphate thus should not be considered alone, as the type of chemical group that decorates these phosphate groups also affects how the membrane reacts to global calcium removal.

We noticed that divalent ions removal increases water penetration in the membrane. It is interesting to note that earlier studies^13,55^ have pointed out that the presence of divalent cations, either Mg^2+^ or Ca^2+^, diminishes water penetration through LPS layers. This increase correlates with the presence of larger shallow and deep defects in the membrane and with the increase in the area per lipid of LPS (see Figure S 7 – S 8).

The reaction of polymyxin-sensitive models, P2 and P1, to such global stress is in line with the conclusions made by Lam and coworkers^13^. In their theoretical work, the authors quantify the electrostatic modifications of the LPS layer using a very coarse-grained model, demonstrating that the presence of divalent ions (Mg^2+^) which induce a negative lateral pressure, which tightens the LPS layer, whereas monovalent ions cause a positive lateral pressure, leading to the swelling of lipid headgroups. In other words, the presence of magnesium ions turns the repulsive interactions between LPS molecules into attractive interactions.

Rice and coworkers made a seminal work regarding the influence of both ion content and phosphate groups in outer membranes models, relying on CHARMM36 force field^45^. They showed that modifications of the lipid A phosphate groups that are linked to resistance mechanisms to polymyxins make the model membranes much less sensitive to the type of ions (monovalent or divalent) used in their simulations. These conclusions are in line with our observations based on the lipid packing defect and the area per lipid.

### Local ion displacement is more energetically costly in polymyxin-resistant membranes

The collective variable we introduce in this work allows us to address how membrane-bound divalent ion displacement is affected by LPS content of the membrane models. Jefferies *et al*.^47^ reported that polymyxins did not induce major changes of the physical properties of the model membranes. Fu and coworkers^41^ also pointed that changes of the lipid packing defect is not significant for polymyxin/lipid ratio under 2 %. Hence, we expected that a local stress that mimics only a portion of polymyxins interactions will not have significant effects on membrane properties, but that ion displacement is likely to be affected by the composition of membranes.

Santos *et al*., by measuring calcium diffusion coefficient in their MD simulations, demonstrate that ion mobility is less affected in polymyxin-resistant model membranes, strongly corroborating our conclusions from free energy calculations^40^. Additionally, they indicate that ion displacement upon polymyxin exposure primarily occurs along the membrane surface, which our simulations support by showing minimal desorption – *i.e*. 1-2 events when displacing all ions from the cylindrical area where we apply the biasing potential – of calcium ions from the membrane (see movie S 1).

It is important to point out that the collective motion calcium-phosphorus might depend on the precise LPS parameterization. Indeed, Rice *et al*.^45^ reported that CHARMM force field tends to overestimate cation binding. They hypothesized that this comes partly from the -2 net charge attributed to the phosphate groups of lipid A. Based on this report, we also simulated the system P1 with -1 net charge for the lipid A phosphate groups which keeps showing that collective motion.

### Polymyxins facilitate ion displacement and ion displacement facilitates polymyxin adsorption

Our results support that an adsorbed polymyxin molecule can increase the mobility of membrane-bound divalent ions. That observation is in agreement with the results obtained by Santos and coworkers^40^, where they led united atom simulations showing that the presence of polymyxin B increases the calcium diffusion coefficient by about a 2-fold rate or higher when the drug is adsorbed on the membrane surface.

The increased ion mobility suggests that polymyxins contribute to ion displacement, even at the early stages of polymyxin-membrane interaction, when the drug is merely adsorbed on the membrane. This aligns with observations by Berglund *et al*. who reported intermittent but no long-term displacement of membrane-bound ions upon polymyxin B_1_ interaction^27^. Our free energy profiles show that while the equilibrium value 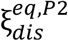 is marginally affected polymyxin presence, the decrease of k_dis_ indicates enhanced calcium ion mobility. Additionally, Jiang *et al*.^43^ report coarse-grained MD simulations showing that divalent ion displacement remains energetically unfavorable in the presence of polymyxin.

It is not clear how polymyxins reach a state upon which they displace membrane-bound divalent cations. Polymyxin insertion remained shallow throughout our simulations, with polymyxins entering no farther than the outer core of LPS. This is in line with previously reported presence of multiple energy barriers for deeper insertion of polymyxin or other relevant drugs^47,54,61–63^. Hence, it is an open question whether the presence of a defect should pre-exist to promote this (partial) translocation of polymyxins or if a concentration threshold should be reached on the membrane to facilitate this permeation, questioning the initial steps of the self-promoted model presented by Hancock^64^. We observe that the presence of a defect in the distribution of divalent ions favorizes the insertion of a polymyxin molecule. Polymyxin-induced defects thus could be self-fed and drive more polymyxin molecules to insert. This observation aligns with numerous studies indicating that polymyxins can penetrate deeply into the hydrophobic region of the outer membrane and even reach the inner membrane^21,23,65^. Particularly, there are experimental evidences that polymyxins accumulate in the KDO / head group region^29^. However, one cannot assume that polymyxin partial translocation is a necessary step in its bactericidal activity, as there are evidences showing that polymyxin internalization across the outer membrane is not necessary to alter OM properties and damage the cell envelope^39,66^.

### Coarse-grained simulations and local stress

The results from our coarse-grained MD simulations using Martini 2 force field^17^ present a notable divergence from the all-atom simulations when analyzing the behavior of divalent ions in the outer membrane. While atomistic simulations showed a widened free energy profile along the collective variable in the presence of polymyxins, indicative of increased ion mobility, the coarse-grained simulations exhibited stiffer free energy profile, suggesting reduced ion mobility, in the presence of polymyxins. This discrepancy can be attributed to the *freezing* effect observed in the coarse-grained simulations, where divalent ions induce localized immobilization of adjacent beads, including phosphate and water molecules.

A recent experimental study highlighted the formation of crystalline structures in lipopolysaccharides (LPS) upon interaction with polymyxins, providing context for our observations^59^. It raises the critical question of whether this freezing is a force field-linked artifact, especially since such freezing was not observed in our atomistic simulations. The free energy profiles, which differ significantly between the simulations with and without polymyxin, underscore the impact of this freezing phenomenon.

The parameterization of divalent ions within Martini 2 force field might be a major contribution to the observed differences with our atomistic simulations. Previous work by Jefferies *et al*.^47^ reported that polymyxin B_1_ could induce local transitions, driving model membranes towards glass-like properties in simulations using Martini 2. Our observations align with these findings, as the collective variable displacing calcium ions— mimicking polymyxin action—drives the membrane to a frozen state. Increasing polymyxin concentration accelerates this transition, corroborating earlier reports. The freezing events are present even in the early steps of our coarse-grained simulations and thus the limited timescales in all-atom simulation are likely not the issue.

## CONCLUSIONS

Over the years, the global picture of polymyxin-mediated bacterial death shows to be intrinsically linked to actions at the outer membrane level. Despite new evidence highlighting the role of porins in the mode of action of polymyxins, direct interactions with lipopolysaccharides remain critical, as they solely explain the mechanisms of some of the polymyxin-resistant mutations. Our work demonstrates that both structural modifications of the outer membrane and dynamical properties of the divalent ions are affected by stresses mimicking parts of polymyxins interactions with the outer membrane.

We showed that polymyxin-sensitive model membranes P2 and P1 are significantly more affected by ions removal than polymyxin-resistant ones, with Ara-4N being the least affected, and PEtN being intermediate. Specifically, P2 exhibits the most significant widening of the LPS area per lipid, the largest increase in the lipid packing defect constant, and the highest proportion of LPS flipping from the outer leaflet to the inner leaflet.

By introducing a new collective variable to monitor the local process of ion displacement in the membrane, we demonstrated that ion mobility is influenced by the type of LPS present. Notably, P2 is more affected than P1, which in turn is more affected than PEtN, with Ara-4N being the least affected. The addition of a single polymyxin molecule further increased ion mobility in the P2 system. Concurrently, we showed that defects in the membrane-bound divalent ion network facilitate the internalization of adsorbed polymyxin, supporting the self-promoted uptake model proposed by Hancock.

Our results also indicate that the employed force field in simulations plays a crucial role in the observed effects of ion displacement on membrane properties. The use of Martini parameterization leads to the freezing of the entire region surrounding the LPS phosphate groups in our coarse-grained systems, which could correspond to experimental observations of LPS crystallization and findings from other studies using this force field. Nevertheless, this transition was not captured in our atomistic simulations and heavily relies on electrostatic interactions with divalent ions.

## METHODS

### Modeling the lipid matrix of the outer membrane of Gram-negative bacteria

Lipopolysaccharides consist of numerous fatty acid chains, typically ranging from 4 to 7. The higher number of tails in lipid A than in most phospholipids confers a higher contact surface between LPS and leads to tighter tails packing^67^. Both properties contribute to the high stability and low fluidity of the OM and to antibiotic resistance. Above lipid A, core oligosaccharides greatly participate in the lamellar structure of the OM. This contribution is attributed to negatively charged moieties, *i.e*. phosphoryl and carboxyl groups, bridged between LPS by a network of divalent ions (Ca^2+^ and Mg^2+^) which further enhances membrane rigidity and stability^7,10,68,69^. The O-antigen, a polymeric chain of saccharides of variable length^16,70^, is covalently bound to the core. Together, they constitute the hydrophilic layer of the OM that favorably interacts with the surrounding medium. In Gram-negative bacteria, LPS exhibit diversity in the number of fatty acid chains, their lengths, and the composition of saccharides in the core and O-antigen. This array of variations defines distinct chemotypes.

Few numerical studies^17,46^, especially at the atomic scale^7,55,71^, utilize smooth LPS, *i.e*. LPS molecule that include the O-antigen. The length of the O-antigen significantly increases the size of the systems, adding substantial challenges to the already long sampling times required for outer membrane models due to slow motions of lipopolysaccharides. Hence, many outer membrane models rely on truncated versions of smooth LPS which conserve the essential parts of LPS that interact with many antibiotic families. A few models represent the outer membrane with LPS modeled solely as lipid A^27,40,72–75^. However, it is common to use deep rough LPS (Re-LPS)^27,40,41,47,62,76,77^, which consist of lipid A bound to the Kdo units of the inner core and which represents the shortest core configuration able to maintain bacterial membrane integrity. These mutants lack core phosphate groups believed to play crucial roles in membrane structure and in its interaction with polymyxins. Therefore, rough LPS should be considered in models focusing on such interactions^78^. Rc-LPS^45,67^ and Ra-LPS^43^ are two commonly described rough LPS, with Rc-LPS being slightly shorter than Ra-LPS. Studies have shown that these additional saccharides influence membrane properties, including the ordering of the phospholipid leaflet^79^ which motivated our choice to rely on Ra-LPS for the outer leaflet of our outer membrane models.

In our work, we focus on models of the outer membrane of *Salmonella enterica*, a pathogen known for causing salmonellosis. *S. enterica* was also shown^80^ to develop resistance against polymyxins relying on modifications of the lipid A phosphate groups that are present in CHARMM-GUI membrane builder^81,82^ online tool, allowing a direct comparison between polymyxin-sensitive and polymyxin-resistant models.

### All-atom models of outer membrane of Gram-negative bacteria

Four models of LPS molecules were used to create four different model membranes (see **Figure 2**). Two models, P2 and P1, represent membranes of non-resistant strains of *Salmonella enterica*, while the two other models represent membranes with PEtN or Ara-4N groups decorating both lipid A phosphate groups. Each membrane is made of 79 LPS molecules in the upper leaflet and 255 POPE molecules in the lower leaflet, so that the area of each membrane shall match one another. One extra system, embedding one polymyxin was also modeled, using P2 system. All-atom MD simulations comprise: 1. an equilibration step, based on the steps proposed by CHARMM-GUI for membrane equilibration, 2. an extra equilibration step in case polymyxin is added to the system, 3. a set of 4 replicas of simulations with calcium ions removal, 4. a steered MD simulation pulling along the collective variable which uses the last frame of the equilibration run, and 5. an umbrella procedure that was used to calculate the free energy profiles using the weighted histogram analysis method^83^ (WHAM).

#### 1. Equilibrium step

To check that our model membranes were at equilibrium, we monitored the changes in box size, as well as the temperature and continued our simulation for more than 100 ns, reaching 300 ns in total.

#### 2. Extra equilibration step in systems containing polymyxin

Simulations with single polymyxin were prepared from the frame corresponding to a simulation time of 100 ns, for the window with the biasing potential applied the highest value used for systems without polymyxin. The system was equilibrated with that single polymyxin, with addition of extra counter-ions to neutralize it, for an extra 100 ns. Then, the same procedures of steered MD simulation and umbrella sampling were applied.

In simulations that involve polymyxin, we decided to mimic the hypothesized aggregation process that supposedly occurs at the position of the defect. To do so, we limited polymyxin only in a concentric cylinder to that of the CV, defined in the same way, but with r_0_=3.0 nm, and an additional upper wall potential distant by 6.0 nm from the plane of the membrane center of geometry that is defined by the lipid tails. That latter potential was present to avoid polymyxin leaving the LPS-containing leaflet during simulations, while allowing polymyxin molecule(s) to rearrange structurally. Both extra potentials were set with a weak potential with a force constant κ = 100 kJ mol^-1^ atom^-1^.

#### 3. Ions removal simulations

Each simulation run involving ions removal was run for 300 ns. Each simulation was started from a random frame from the last 100 ns of the equilibrated simulations. Solvent, as well as all membrane-bound ions were removed, and the system was re-solvated with as many chloride ions as in the original system and neutralized with addition of sodium ions. The box size was increased to 20 nm before equilibration along z-axis, which corresponds to an increase in the number of water molecules by 45 to 65 %, depending on the system. This was done to ensure that fluctuations of the membrane while out-of-equilibrium will not drive the system to self-interact. For these systems, we also monitored the changes in the density profiles of representative groups and waited until the two last 25 ns blocks of simulations show no variations that would be bigger than the statistical fluctuations expected in our systems. Hence, we could consider the last two blocks of our whole trajectory, *i.e*. the last 50 ns, to be close to equilibrium. The last 25 ns were used for

Table **3**. Errors on the free energy calculations were calculated by means of bootstrapping method using 200 bootstraps. analysis. Four independent replicas were always used to obtain the averages

**Table 3.** Simulation times for the production runs of the different systems. Note that the equilibration time with polymyxin molecule (P2-PME system) started after inserting the polymyxin on the equilibrated P2 membrane.

| System | Equilibration | Steered MD | Umbrella sampling | Number of windows |
| --- | --- | --- | --- | --- |
| P2 | 300 ns | >75 ns | 300 ns / window | 77 |
| P1 | 300 ns | >75 ns | 200 ns / window | 21 |
| PEtN | 300 ns | >75 ns | 200 ns / window | 29 |
| Ara-4N | 300 ns | >75 ns | 200 ns / window | 29 |
| P2-PME | 100 ns | >75 ns | 200 ns / window | 68 |

#### 4. Steered MD simulations

Our steered MD simulations consisted of a run with a rate of ∼1.1 ions / nm^2^ / ns or 5 ions / ns and a force constant κ = 2000 kJ mol^-1^ ion^-1^. We monitored the reaction of the system to the applied bias by plotting the value of the measured CV versus the value of the applied bias (see Figure S 17). This setting results in a linear response for which the shift between the measured value and the applied bias is always very small, the slope of the linear regression being 1.02 for a correlation coefficient of 0.997.

#### 5. Umbrella sampling procedure

The umbrella procedure was carried by applying a biasing potential with a force constant κ = 500 kJ mol^-1^ ion^-1^. Figure S 18 shows that the histograms had enough overlaps. We also checked the convergence of the free energy by analyzing 25 ns blocks of our simulations (see figure S 19). Simulation times, number of windows, and equilibration times are reported in

### Definition of the collective variable

The collective variable is defined as the number of calcium ions in a cylinder of r_0_=1.2 nm of radius centered in the center of geometry of the membrane without periodic images. The radius of that cylinder was chosen to encompass the characteristic size of an aggregate of polymyxins, noting that a growing number of observations drive to the conclusion that polymyxins do not access the membrane surface individually^33,27,41^. Soft boundary conditions defined by a switching function such that 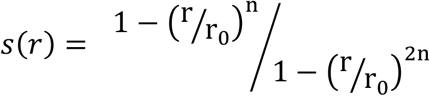, where n=14, were applied.

As visible in the movie S1, most degrees of freedom of the calcium ions are not affected by our CV, resulting in ions that generally move radially, due to the lamellar substructure of the LPS, but also to few events of ions exiting the membranes (in orange in the movie).

Our collective variable is defined as a number of calcium ions in PLUMED library^84^. Values are then converted, in order to express our collective variable as a surface density, by straightforwardly dividing the values of the collective variable by the surface of the cylinder, *i.e*. by *1.2*^*2*^ *π*.

Unless stated otherwise, our collective variable was ported to coarse-grained system without modifications.

### Parameters used with atomistic simulations

All atomistic simulations that required an equilibration procedure used a set of energy minimization, canonical ensemble simulation, and isobaric-isothermal ensemble. These simulations gradually alleviate a set of position restraints that affects the lipids, while increasing the time step (in case of NVT and NpT runs). The NVT and NpT runs are relying on Berendsen algorithm for the thermostat and/or the barostat for the first short steps, according to the protocol provided by CHARMM-GUI^82^, before switching to Nosé-Hoover algorithm for the thermostat, and Parrinello-Rahman algorithm for the barostat, using a coupling constant of 1.0 ps and 5.0 ps, respectively. Temperature coupling was made separately for the membrane and the solvent. The simulations are held at 310 K and 1.0 bar. Dispersion interactions are treated with a cut-off set at 1.2 nm where forces are smoothly switched to 0 from 1.0 nm. Electrostatic interactions are treated using the fast smooth particle-mesh Ewald^85^ (SPME) algorithm, implemented in Gromacs package version 2022.3^86^ that was used all along this work and patched with PLUMED library^84^ version 2.8.1^87^. All simulations used for producing the data that was later analyzed used a 2 fs time step with all hydrogen bonds treated as constraints using LINCS algorithm^88^.

### Coarse-grained models of outer membrane of Gram-negative bacteria

Coarse-grained models are all relying on the Martini equivalent of our P2 model. We used the default Ra LPS available from CHARMM-GUI^82^ and built an asymmetric membrane with 81 LPS on the outer leaflet and 246 POPE in the inner leaflet. Three systems were modeled for energy calculation and a fourth (0, 1, and 5 polymyxins).

Each of the three systems was first equilibrated, before proceeding to a steered MD procedure and energy calculations by means of umbrella sampling and WHAM^83^. Equilibration was led following CHARMM-GUI^82^ protocol adapted to Martini force field^89^ (similar to the one for CHARMM force field^44^ described above) and terminated by a 2 µs long equilibration run. The steered MD simulation consisted of a 10 ns relaxation with no bias applied, followed by an additional 10 ns at close to the equilibrium value (15 ions in the cylinder). The run to reach a minimal value along the CV was of 1 microsecond. Then umbrella sampling procedure was applied for all the systems between 0 and 20 ions in the cylinder. Simulations times are reported in Table **4**.

**Table 4.** Simulation times for the production runs of the different systems. Note that the equilibration time with polymyxin molecules of 1 µs comes after inserting the drugs on the equilibrated membrane.

| Number of polymyxins | Equilibration | Steered MD | Umbrella sampling | Number of windows |
| --- | --- | --- | --- | --- |
| 0 | 2 μs | >1 μs | 3 μs / window | 73 |
| 1 | 1 μs | >1 μs | 2 μs / window | 72 |
| 5 | 1 μs | >1 μs | 3 μs / window | 81 |

### Analyses

We estimated the defect size constant using Fatslim^90^ and used the 3^rd^ carbon that branches the 5^th^ lipid and 6^th^ lipid tails to differentiate deep from shallow defects. We consider here that we are interested in deep defects that are buried in the fatty acid medium and chose this carbon as it is related to a tail that is central in our LPS model.

To compute the proportion of LPS flipping, the density profile was integrated. The density is centered in 0, which corresponds to the center of geometry of the methyl groups of LPS and POPE. The proportion of flipped LPS is the integral of all values lower than 0 divided by the integral of the whole profile. Only the last block of trajectory was used (25 ns) for each replicate. Errors are computed as the standard error over the four replicas.

## Supporting information

Movie showing a steered MD simulation illustrating the collective variable introduced in the manuscript

Movie showing a steered MD simulation where collective motion between calcium ions and phosphorus atoms is visible

Supplemental figures to the manuscript

## ASSOCIATED CONTENT

### Data and Software availability

Initial configurations, input parameters, and topologies are available at https://zenodo.org/records/13486348.

### Supporting information

The Supporting information is available free of charge.

The analysis of the density profiles of relevant groups for the global stress, to the structure of the lipid membrane, when an equilibrium is reached, the depth at which water molecules penetrate the LPS leaflet, the distribution of the area per lipid of LPS molecules, the data used to compute the percentage of LPS flipping, the defect size constant for the local stress, the distribution of the minimal distance between phosphorus atoms of LPS molecules, the orientation of LPS molecule when a local stress is applied, the RMSF of lipid A phosphorus atoms when a local stress is applied, the measurement of the reaction of the system to the moving biasing potential during the steered MD simulation, and the histograms associated with the umbrella sampling procedure are available in one file (PDF).

A movie of the steered MD simulation for P2 illustrating the collective variable (MP4).

A movie of the steered MD simulation for P2 showing the collective motion of phosphorus atoms with calcium ions, and the absence of such co-motion for the aliphatic carbons (MP4).

## Author Contributions

M.S. and T.R. designed and carried out the simulations and the analysis. T.R. supervised the research. All the authors contributed to the discussion, writing, and revising of the manuscript.

## ACKNOWLEDGMENT

The work was supported by the European Regional Development Fund-Project " MSCAfellow5_MUNI “(No CZ.02.01.01/00/22_010/0003229), and by the project National Institute of virology and bacteriology (Programme EXCELES, ID Project No. LX22NPO5103) - Funded by the European Union - Next Generation EU. We acknowledge VSB – Technical University of Ostrava, IT4Innovations National Supercomputing Center, Czech Republic, for awarding this project access to the LUMI supercomputer, owned by the EuroHPC Joint Undertaking, hosted by CSC (Finland) and the LUMI consortium through the Ministry of Education, Youth and Sports of the Czech Republic through the e-INFRA CZ (grant ID: 90254). Part of this work was performed using computing resources of CRIANN (Normandy, France).

## ABBREVIATIONS

MDR: multidrug-resistant
IM: inner membrane
OM: outer membrane
PE: phosphatidylethanolamine
PG: phosphatidylglycerol
CL: cardiolipins
LPS: lipopolysaccharides
DAB: L-diaminobutyric acid
PMB: polymyxin B
PME: polymyxin E, colistin
MD: molecular dynamics
PEtN: phosphoethanolamine (PEtN)
Ara-4N: 4-aminoarabinose
GalN: galactosamine
EDTA: ethylenediaminetetraacetic
LOESS: local regression smoothing
RMSF: root-mean-squared fluctuations
WHAM: weighted histogram analysis method
SPME: smooth particle-mesh Ewald

## Notes

https://zenodo.org/doi/10.5281/zenodo.13486347

### Competing Interest Statement

The authors have declared no competing interest.

