## Supplemental figures to the manuscript for "Role of Divalent Ions in Membrane Models of Polymyxin-Sensitive and Resistant Gram-Negative Bacteria"

SUPPORTING INFORMATION

### Analyzing the outcome of a global stress to the model membranes

We calculated the free energy profiles of phosphorus atoms in equilibrated model membranes to check for the presence of a defined peak around the lipid A phosphate groups.

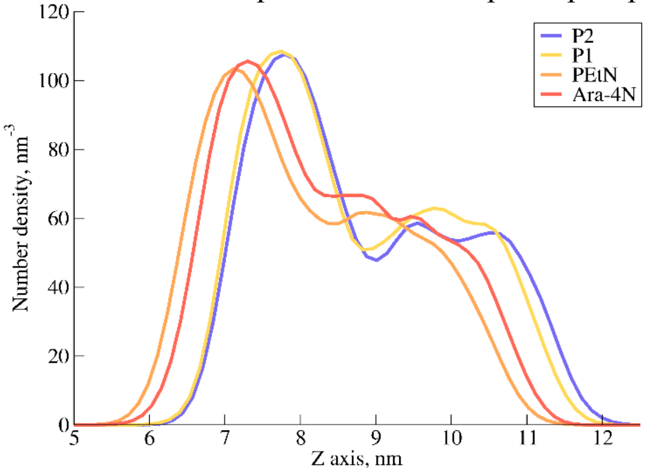


Figure S 1 Density profiles of the phosphorus atoms for the different model membranes used in our all-atom simulations in this work. One can clearly see the presence of a first well defined peak associated with the phosphorus atoms from the phosphate groups present in lipid A head group.

To check our analyzes on the systems under global stress, *i.e.* where all calcium ions have been removed, and replaced by sodium ions that were placed in the solvent, we first computed the density profiles along the normal to the membrane, for each replicate, and for each model membrane (see Figure S 2, Figure S 3, Figure S 4, and Figure S 5). We computed the density profiles of nine distinct groups, namely, LPS tails, lipid A head group, LPS inner core, outer core, POPE tails, head group, sodium ions, chloride ions, and water molecules. Each subplot shows the individual density profiles for each of the four replicates that were produced. Each subplot shows correspond to a chunk of 30 ns of simulation, the starting time being indicated on top of the given column, in nanoseconds. One can see that major rearrangement occurs for all systems, mainly during the first ~200 ns and that the individual profiles of each replicate do not defer drastically for the last two chunks. Finally, we considered that 300 ns of simulation was enough since the two last chunks show profiles that differ only marginally, in a way that can be attributed to normal statistical fluctuations. Since 30 ns of statistics would be insufficient for an ensemble average for our systems, all our analyzes are made on the 4 replicates, giving us the equivalent of 120 ns of equilibrated simulation run.

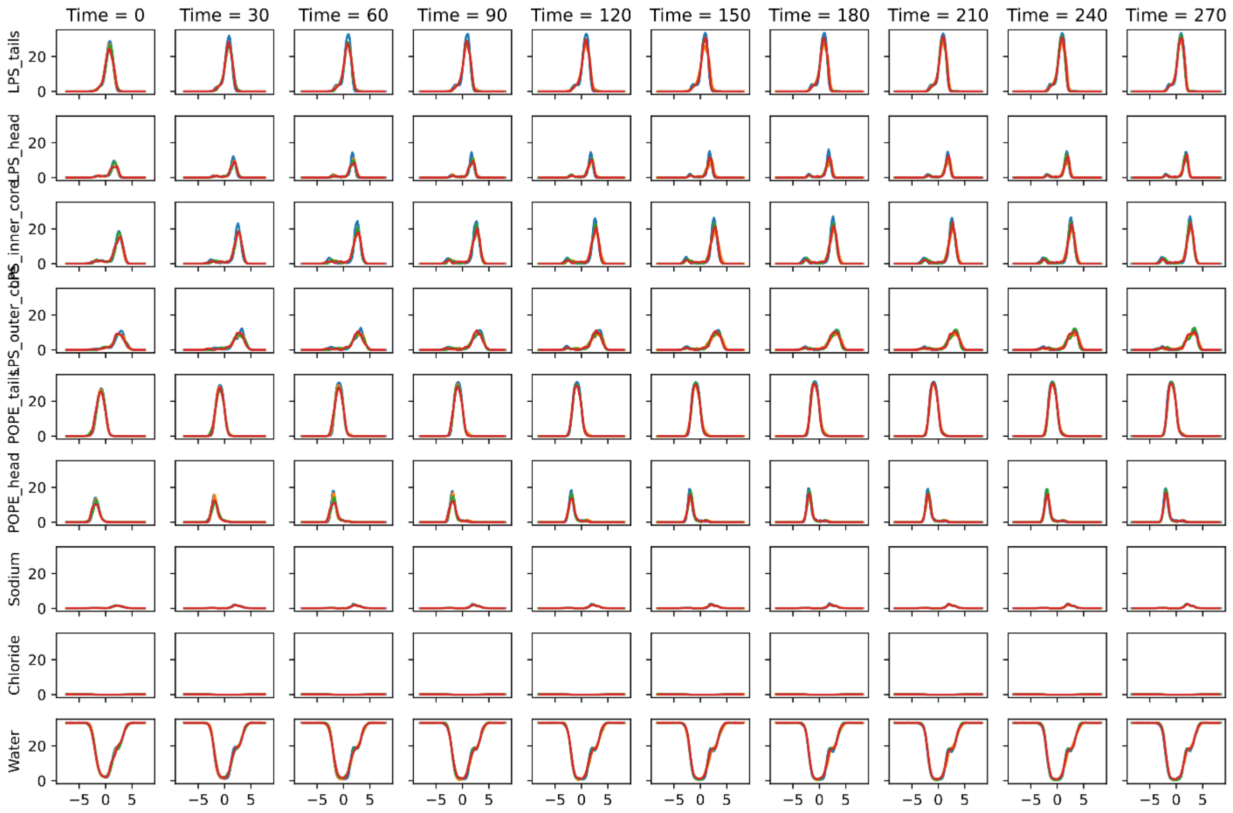


**Figure S 2.** Density profiles for system P2. Densities are number densities, in $nm^{-3}$. The densities are plotted along the normal to the membrane, and the values are expressed in nanometers. The center of the abscissa lies in the center of geometry of the methyl groups from both LPS and POPE molecules. Each subplots displays the four replicates for the given system in red, orange, green, and blue.

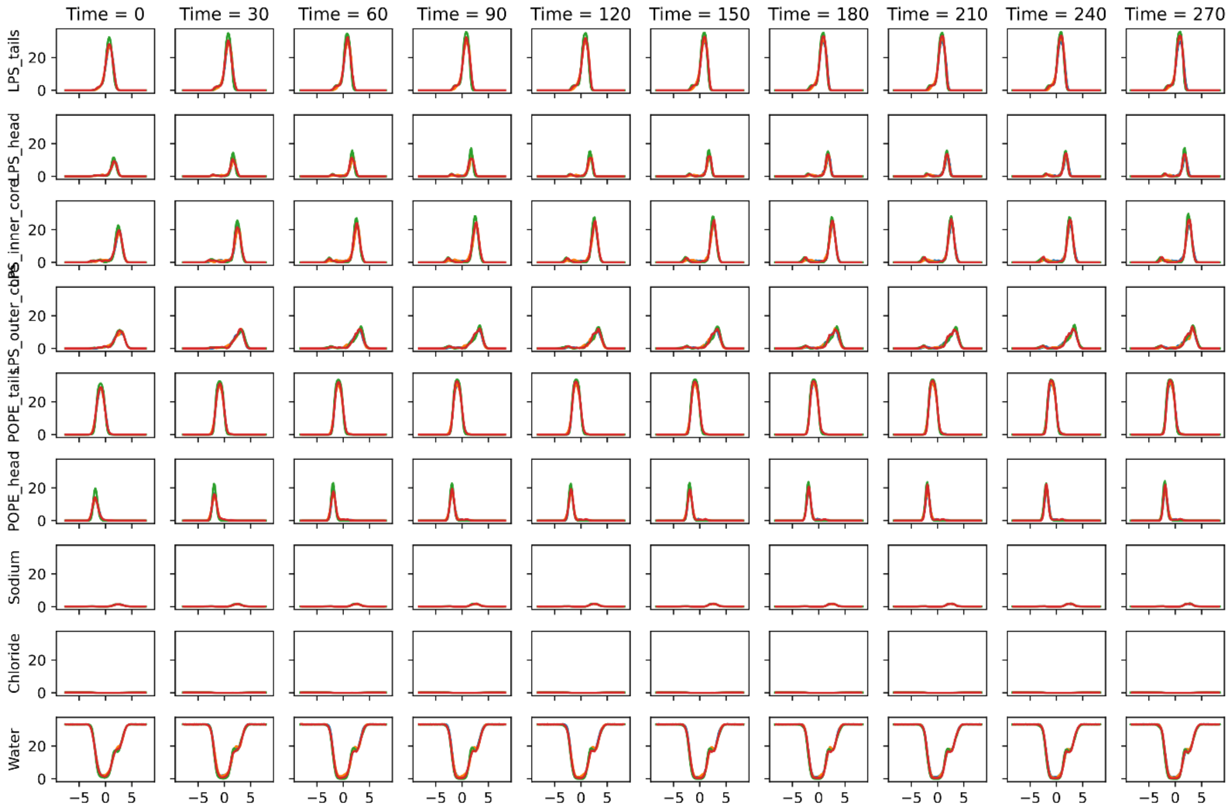


**Figure S 3.** Density profiles for system P1. Plots are made the same way as for system P2.

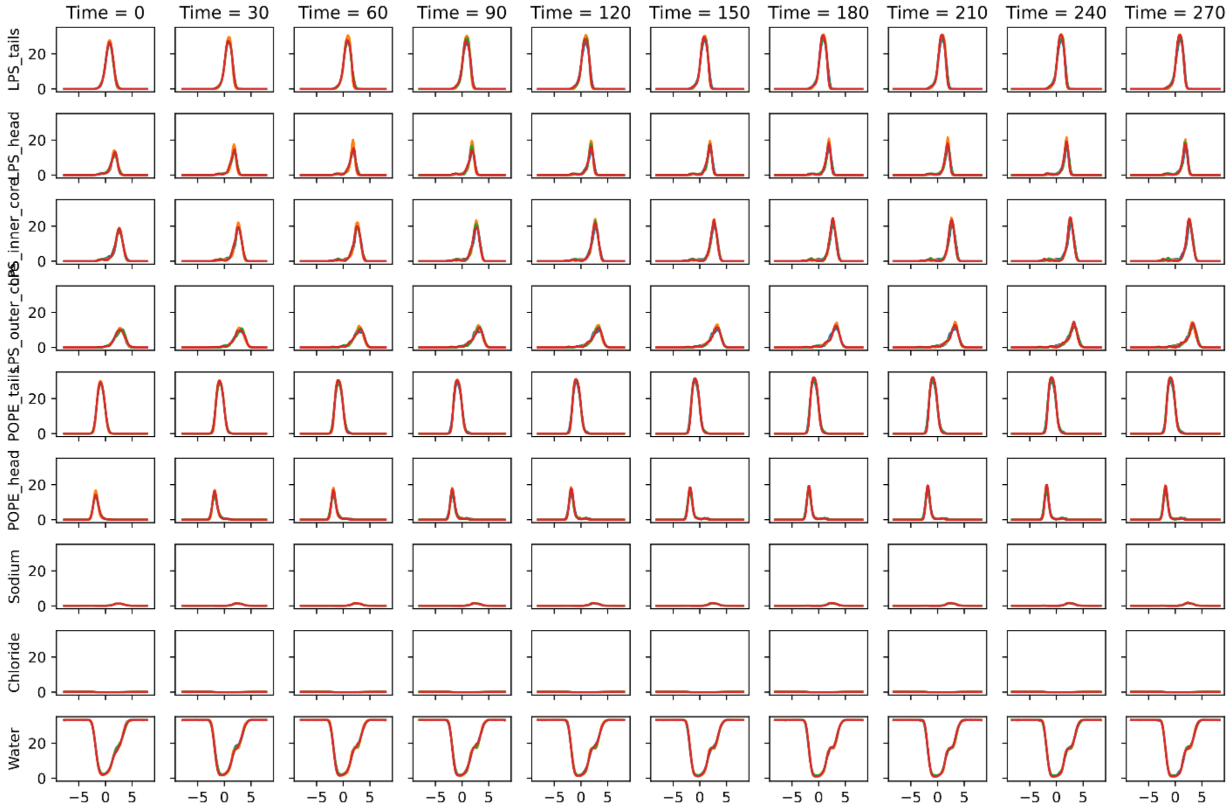


**Figure S 4.** Density profiles for system PEtN. Plots are made the same way as for system P2.

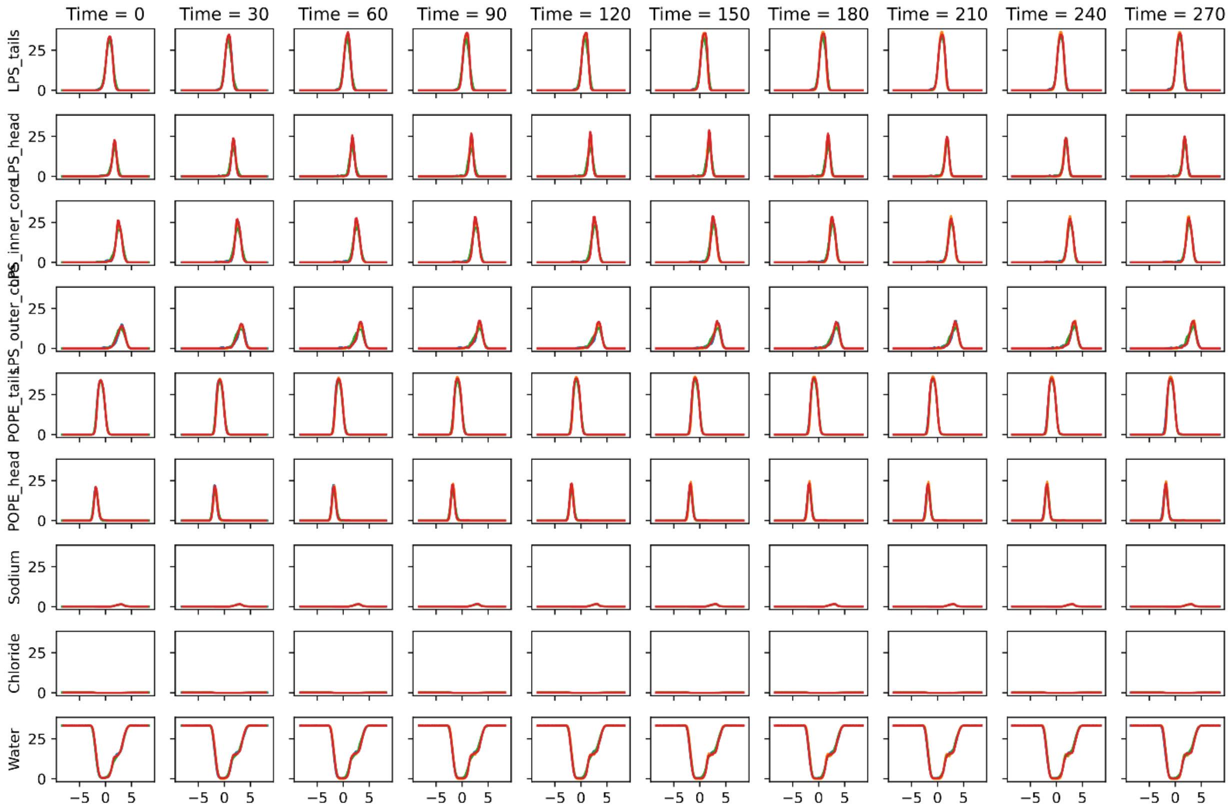


**Figure S 5.** Density profiles for system Ara-4N. Plots are made the same way as for system P2.

In order to estimate the proportion of LPS flipping to POPE leaflet upon calcium ions removal, we have used the averaged density of LPS head groups for each system. For a given system where the density was computed between $z_{min}$ and $z_{max}$, the proportion of flipped LPS, $P_{flipped}$, was estimated as: $P_{flipped} = \int_{z_{min}}^{0} d_{\text{LPS head group}}(z)dz \Big/ \int_{z_{min}}^{z_{max}} d_{\text{LPS head group}}(z)dz$.

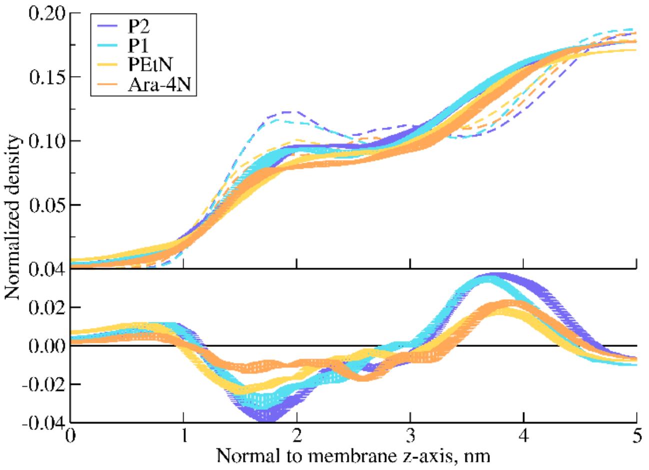


**Figure S 6.** Density profiles of water molecules in the LPS leaflet. Profiles in dashed lines are systems with no stress, while profiles in plain lines are systems under global stress.

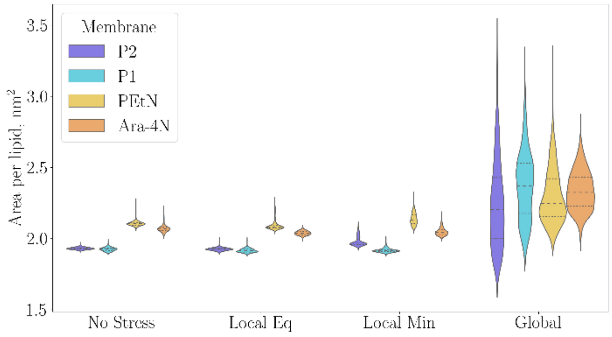


**Figure S 7.** Violin plot of the distribution of the area per lipid for the four model membranes used for all-atom simulations and for the 3 scenarios of stress (no stress, local stress close to the equilibrium value, or minimum value of the collective variable for that stress, and global stress.

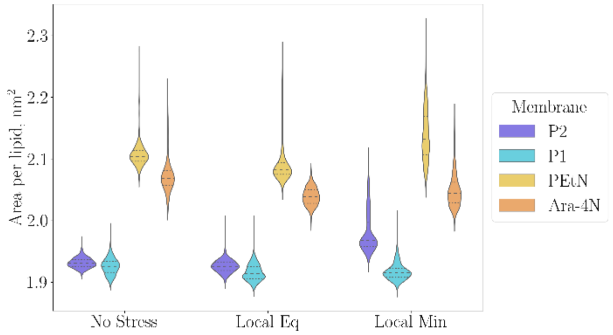


**Figure S 8.** Violin plot of the area per lipid showing only the membranes at equilibrium (no stress) and with the local stress (with the collective variable introduced in our work).

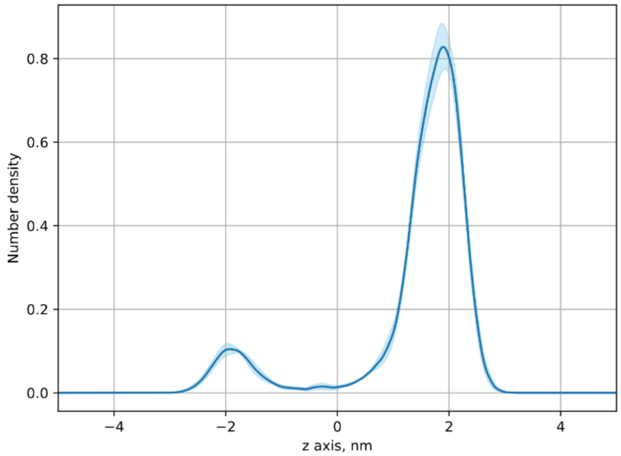


**Figure S 9.** System P2. Averaged density of lipid A head group. The error bar is the standard error computed from the four replicates.

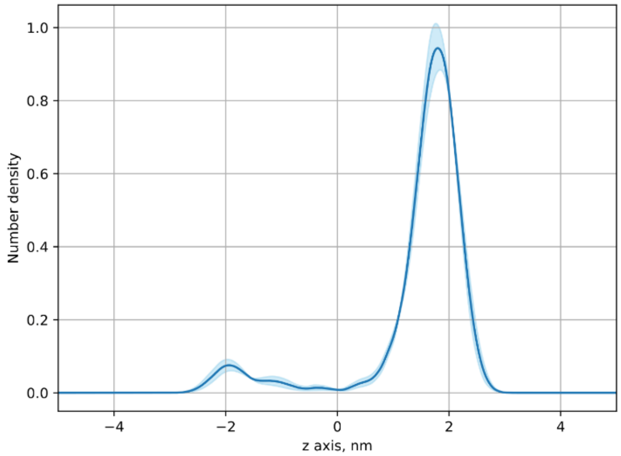


**Figure S 10.** System P1. Averaged density of lipid A head group. The error bar is the standard error computed from the four replicates.

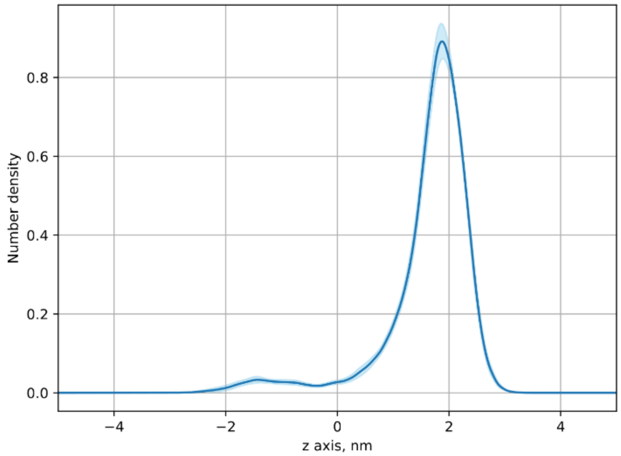


**Figure S 11.** System PEtN. Averaged density of lipid A head group. The error bar is the standard error computed from the four replicates.

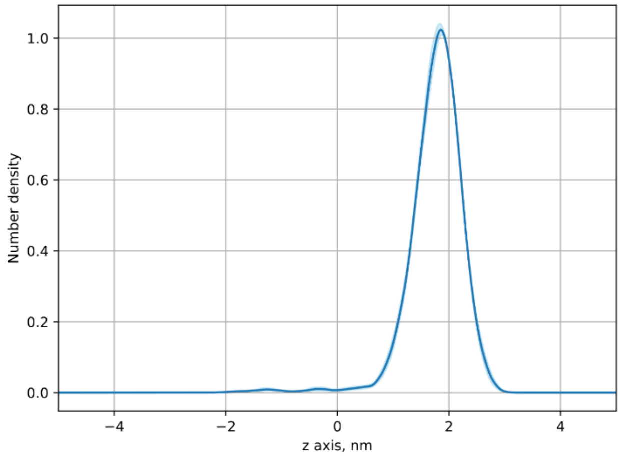


**Figure S 12.** System Ara-4N. Averaged density of lipid A head group. The error bar is the standard error computed from the four replicates.

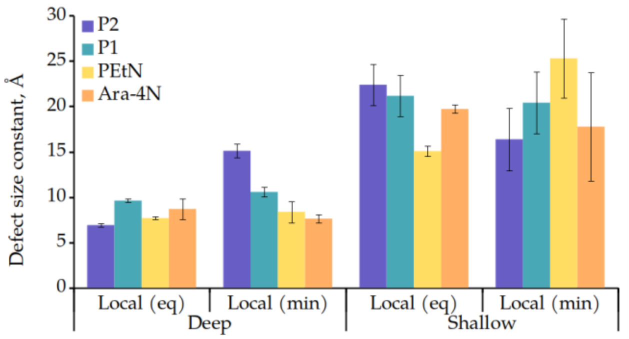


**Figure S 13.** Defect size constant for membrane biased locally with the collective variable presented in the present work. In the figure, (eq) refers to the value of the collective variable held to the closest value of its equilibrium value, and (min) refers to the minimal value reached using this collective variable.

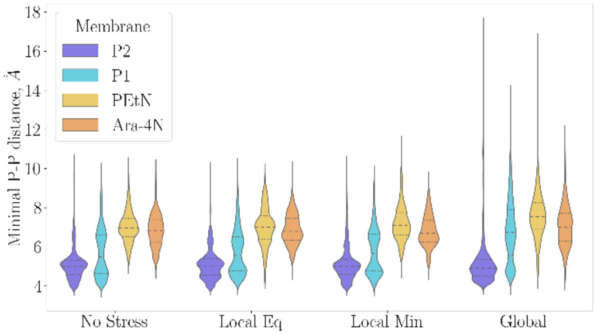


**Figure S 14.** Violin plot of the distribution of the minimal distance between lipid A phosphorus atoms of different LPS molecules for the four model membranes used for all-atom simulations and for the 3 scenarios of stress applied on divalent ions.

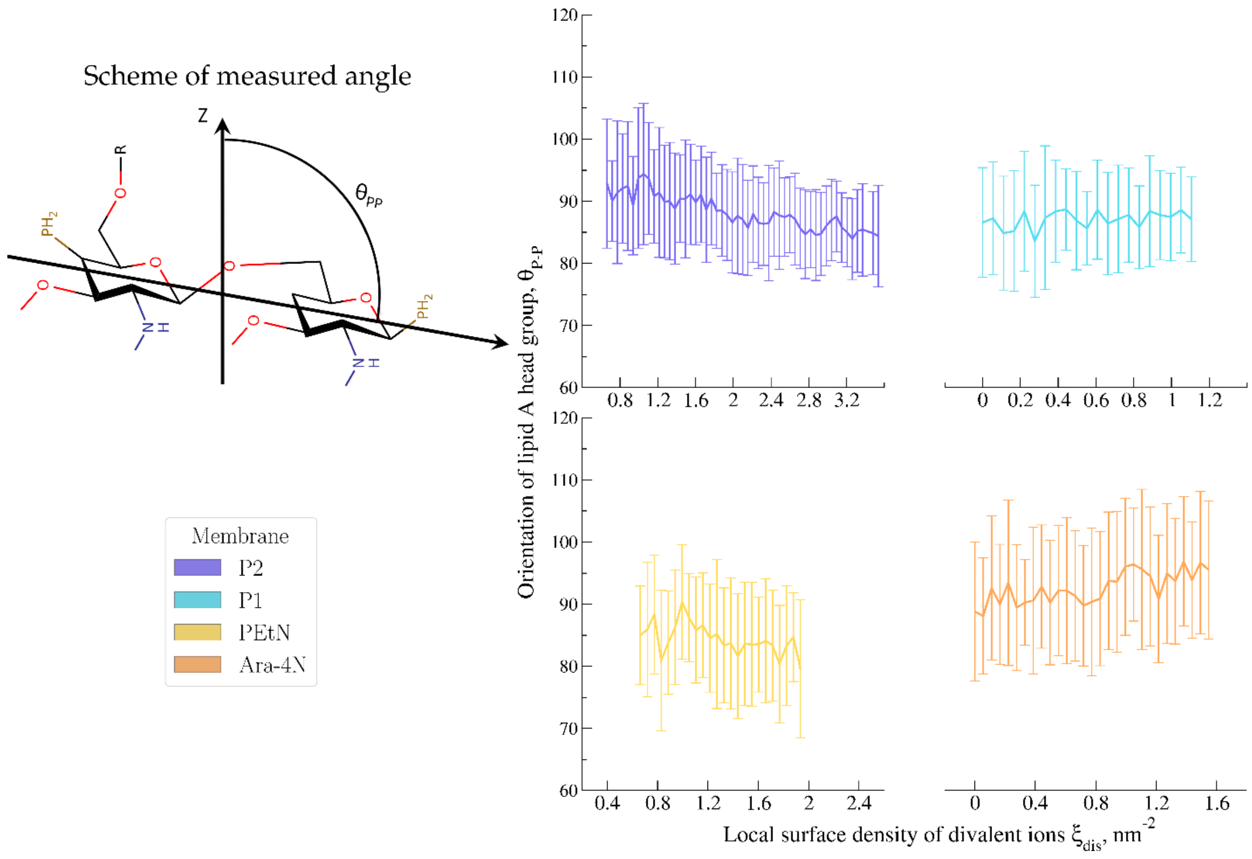


**Figure S 15.** Measure of the angle to the normal of the membrane made by the vector formed by lipid A phosphate atoms, for each of the models used for atomistic simulations.

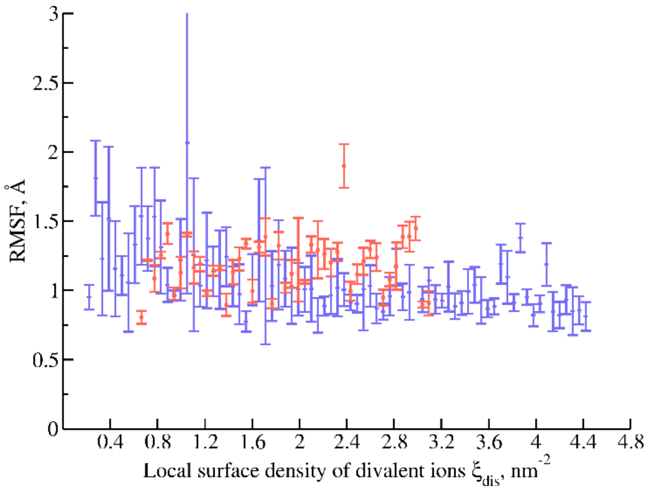


**Figure S 16.** Root mean-squared fluctuations of lipid A phosphorus atoms are monitored for the system P2 without (mauve) and with (vermilion).

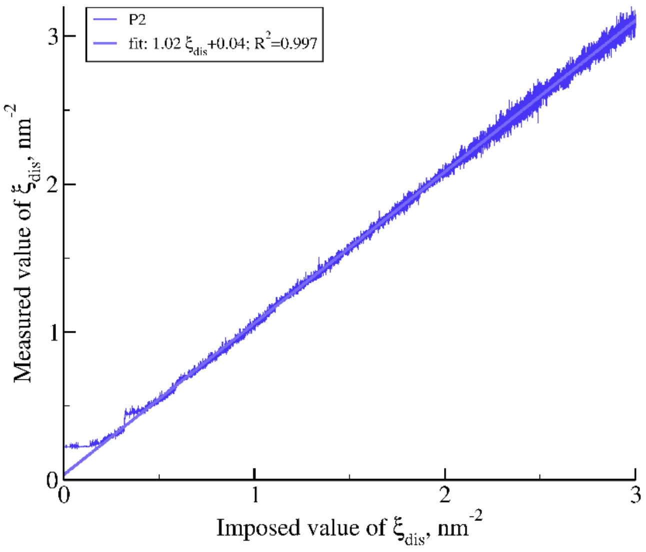


**Figure S 17.** Measurement reaction of the system to the moving biasing potential applied during the steered MD simulation. In addition to direct controls, we monitor the value of the collective variable reached by the system that is biased to check that the response is linear, and that there is no delay to that response. This is an additional control to ensure that the rate chosen for the steered MD procedure is not too fast.

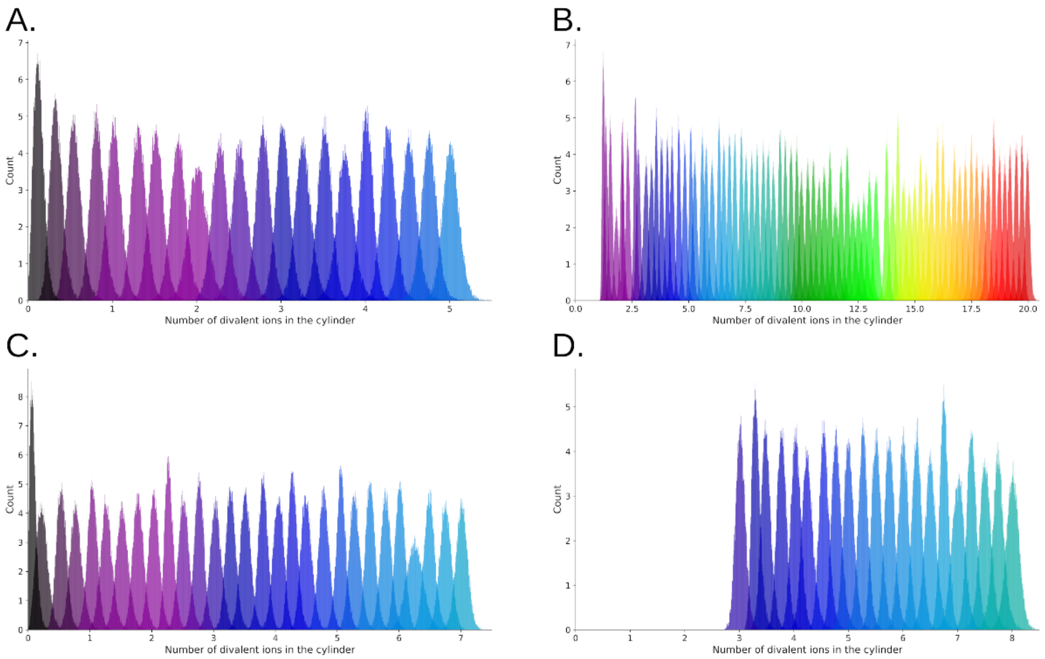


**Figure S 18.** Histograms of the values of the collective variable reached by the different model membranes for the different windows that were sampled during the umbrella sampling procedure. In panel A. system P1 is reported, in B. P2, in C. PEtN, and in D. Ara-4N.

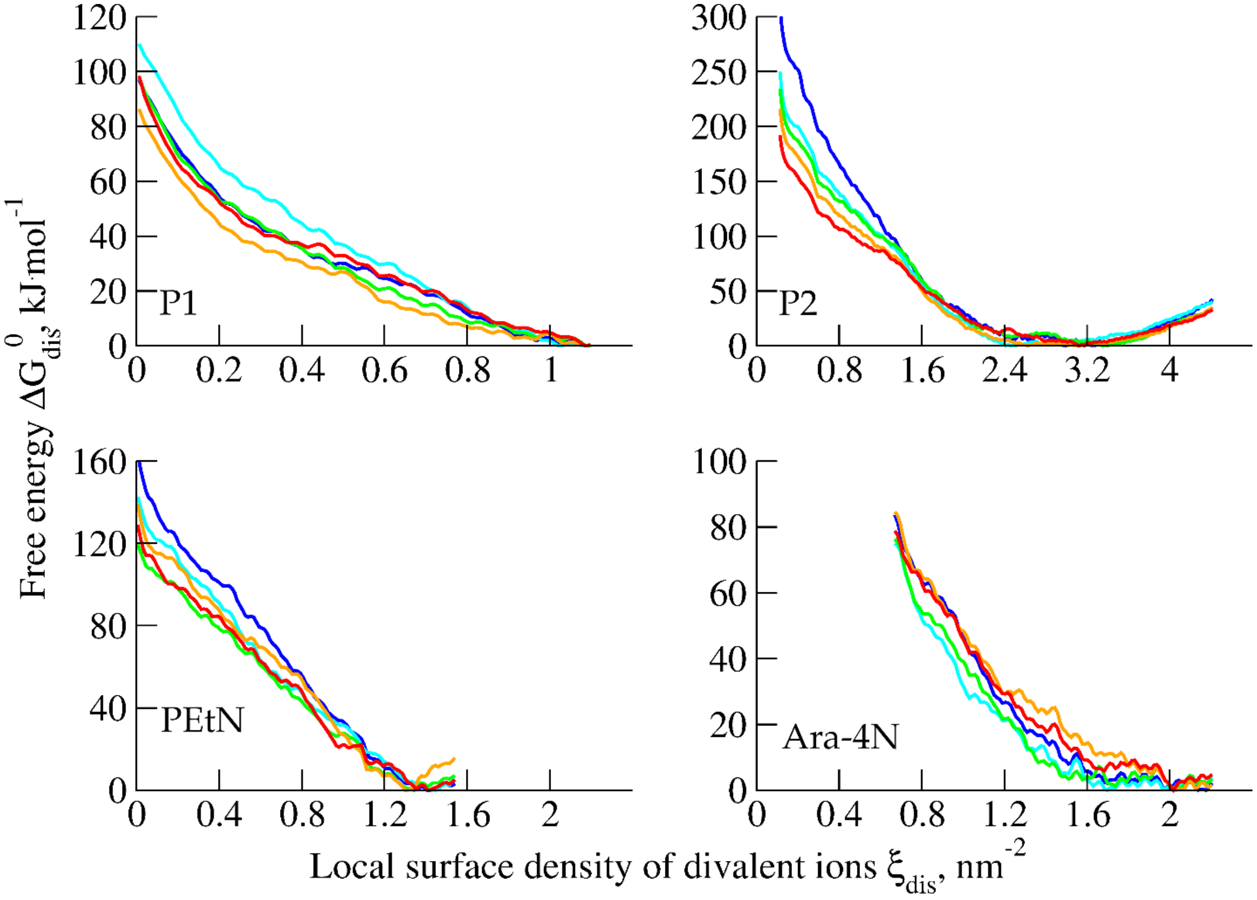


**Figure S 19.** Plots used to monitor the convergence of the free energy calculations. Each line represents 25 ns of simulation per window (P1, PEtN, Ara-4N) or 50 ns per window (P2). Hence, for P1, PEtN, and Ara-4N, the last 125 ns are shown, and for P2, the last 250 ns. The color code is the following, from less sampling to more sampling: dark blue, cyan, green, orange, and red.

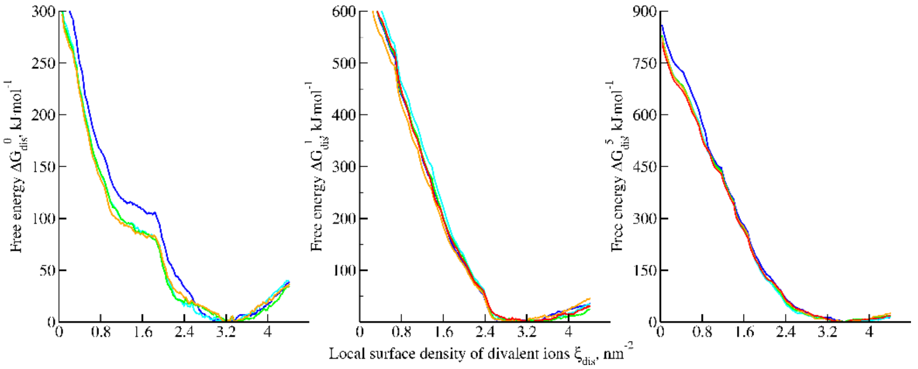


**Figure S 20.** Plots used to monitor the convergence of the free energy calculations. Each line represents 200 ns of simulation per window for systems with 0, 1, and 5 colistin molecules in presence. The color code is the following, from less sampling to more sampling: dark blue, cyan, green, orange, and red.
